# Dendritic spine morphology regulates calcium-dependent synaptic weight change

**DOI:** 10.1101/2021.05.06.442994

**Authors:** M. K. Bell, M. V. Holst, C. T. Lee, P. Rangamani

## Abstract

Dendritic spines act as biochemical computational units and must adapt their responses according to their activation history. Calcium influx acts as the first signaling step during post-synaptic activation and is a determinant of synaptic weight change. Dendritic spines also come in a variety of sizes and shapes. To probe the relationship between calcium dynamics and spine morphology, we used a stochastic reaction-diffusion model of calcium dynamics in idealized and realistic geometries. We show that despite the stochastic nature of the various calcium channels, receptors, and pumps, spine size and shape can modulate calcium dynamics and subsequently synaptic weight updates in a deterministic manner. Through a series of exhaustive simulations, we find that the calcium dynamics and synaptic weight change depend on the volume-to-surface area of the spine. The relationships between calcium dynamics and spine morphology identified in idealized geometries also hold in realistic geometries suggesting that there are geometrically determined deterministic relationships that may modulate synaptic weight change.

## 1 Introduction

Dendritic spines are small protrusions along the dendrites of neurons that compartmentalize post-synaptic biochemical, electrical, and mechanical responses. These subcompartments house the majority of excitatory synapses and are key for neuronal communication and function (*1, 2*). Because of their unique biochemical compartmentation capabilities, spines are thought of as computational units that can modify their synaptic strength through a process called synaptic plasticity (*1, 3*). There have been many different approaches, both experimental and computational, to understand how these small subcompartments of excitatory neurons can regulate learning and memory formation (*4, 5*). These studies have helped identify a few key scientific threads – (a) the biochemical signal transduction cascades in spines spans multiple timescales but Ca^2+^ is the critical initiator of these events; (b) spines have distinct morphological features and these can be categorized depending on physiological or pathological conditions (see Table 1) (*6*); and (c) synaptic weight update is a measure of the strength of a synapse, and represents the strength of the connection between neurons. Synaptic weight represents changes to synaptic connection strength that occurs during synaptic plasticity, such as during long term potentiation (LTP) and long term depression (LTD) (*4*). In this work, we focus on bridging these different ideas by asking the following question – how does spine morphology affect synaptic weight update? To answer this question, we develop a computational model that focuses on the stochastic dynamics of Ca^2+^ in spines of different geometries and map the synaptic weight update to geometric parameters.

**Table 1:**
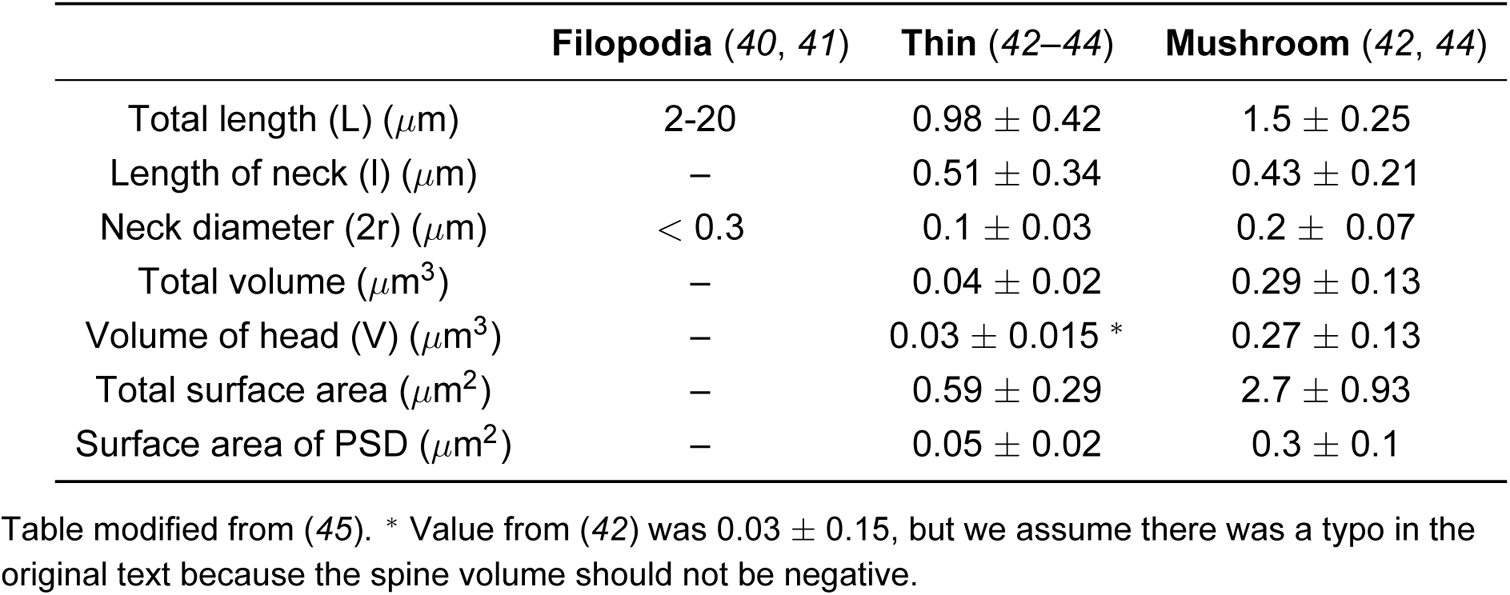
Geometric parameters of different spine morphologies.

|  | **Filopodia (40, 41)** | **Thin (42–44)** | **Mushroom (42, 44)** |
| --- | --- | --- | --- |
| Total length (L) ($\mu\text{m}$) | 2-20 | $0.98 \pm 0.42$ | $1.5 \pm 0.25$ |
| Length of neck (l) ($\mu\text{m}$) | – | $0.51 \pm 0.34$ | $0.43 \pm 0.21$ |
| Neck diameter (2r) ($\mu\text{m}$) | $< 0.3$ | $0.1 \pm 0.03$ | $0.2 \pm 0.07$ |
| Total volume ($\mu\text{m}^3$) | – | $0.04 \pm 0.02$ | $0.29 \pm 0.13$ |
| Volume of head (V) ($\mu\text{m}^3$) | – | $0.03 \pm 0.015^*$ | $0.27 \pm 0.13$ |
| Total surface area ($\mu\text{m}^2$) | – | $0.59 \pm 0.29$ | $2.7 \pm 0.93$ |
| Surface area of PSD ($\mu\text{m}^2$) | – | $0.05 \pm 0.02$ | $0.3 \pm 0.1$ |

Table modified from (45). \* Value from (42) was $0.03 \pm 0.15$ , but we assume there was a typo in the original text because the spine volume should not be negative.

Calcium plays a key role as a second messenger in biochemical and physical modifications during synaptic plasticity, triggering downstream signaling cascades within dendritic spines, and the entire neuron (*3, 7, 8*). Theoretical efforts have linked calcium levels to synaptic plasticity change through a parameter called synaptic weight that essentially captures the strength of the synapse (*9–12*). An increase in synaptic weight is associated with synapse strengthening, while a decrease in synaptic weight is associated with synapse weakening (*13, 14*). While changes in synaptic strength require a host of downstream signaling and mechanical interactions (*15, 16*), the level of calcium is a well-accepted indicator of synaptic plasticity and weight (*10, 17*). This led to the hypothesis that synaptic plasticity outcome could be determined from the calcium dynamics alone; this theory has been readily used for numerous models in computational neuroscience (*10, 18*). Due to their probabilistic nature and discrete number, calcium ion channels and receptors appear to behave stochastically (*19–21*). This indicates that calcium dynamics in the spine leans towards stochasticity and it has been suggested that synaptic plasticity itself relies on stochasticity for robustness (*19, 22–26*). In this work, we seek to understand how spine morphology can modulate synaptic weight update predicted through stochastic calcium dynamics.

Dendritic spines have characteristic sizes and shapes that dynamically change over time in response to stimulus, and are associated with their function and synaptic plasticity (*27, 28*). Just as whole cell shape is known to influence signaling dynamics (*29–34*), studies have specifically probed the interplay between calcium dynamics and dendritic spine morphology (*7, 35–37*). Due to the historical significance of dendritic spines as electrical subcompartments, the morphology of the spine neck has been implicated in regulating calcium signaling and longer spine necks were found to decouple spine-dendrite calcium signaling (*38*). Additional modeling work coupled actinmyosin contractions to cytoplasmic flow to identify two timescales of calcium motion, driven by flow and diffusion respectively, that depend on spine geometry (*39*). A combined analytical and numerical study showed how geometry and curvature gives rise to pseudo-harmonic functions that can predict the locations of maximum and minimum calcium concentration (*36*). More recently, we used a deterministic reaction-diffusion model to investigate dendritic spine morphology and ultrastructure, and found that dendritic spine volume-to-surface area ratios and the presence of spine apparatus modulate calcium levels (*35*). As we have shown before, the natural lengthscale that emerges for reaction-diffusion systems with boundary conditions that have influx and efflux rates is the volume-to-surface area ratio (*29, 36*). What remains unclear is whether the trends from dimensional analysis of deterministic models continue to hold despite the stochastic nature of calcium influx and efflux across the wide range of spine shapes.

In this work, using idealized and realistic spine geometries, we investigate the impact of shape and stochasticity on calcium dynamics and synaptic weight change. We seek to answer the following question: How do specific geometric parameters – namely shape and size of dendritic spines – influence calcium dynamics and therefore synaptic weight change? To address this question, we built a spatial, stochastic model of calcium dynamics in different dendritic spines geometries. We used idealized geometries informed by the literature to control for the different geometric parameters and then extended our calculations to realistic geometries. We probed the influence of spine shape, volume, and volume-to-surface area ratio on calcium influx, variance of calcium dynamics, and the robustness of synaptic weight. We show that although calcium dynamics in individual spines are stochastic, synaptic weight changes proportionally with the volume-to-surface area of the spines, suggesting that there are deterministic relationships between spine morphology and strengthening of synapses.

## 2 Model Development

Ca^2+^ dynamics in dendritic spines have been previously studied using computational models (*35–37, 46, 47*). In this work, we focused on modeling effort on the early, rapid influx on Ca^2+^ for spines of different sizes and shapes with the goal of identifying relationships between spine geometry and early synaptic weight change. Our model is based on previous works (*18, 35, 37*) with some modifications and simplifications to enable us to identify the relationship between spine morphology and synaptic weight change. Inspired by (*37*), we converted a previous deterministic model of calcium influx (Bell *et al.* (*35*)) to a spatial, particle-based stochastic model constructed in MCell (Monte Carlo Cell) (*48–50*), to capture the stochastic nature of Ca^2+^ dynamics in the small spine volumes. We specifically focus on dendritic spine geometries and calcium dynamics representative of hippocampal pyramidal neurons (*35*). We detail the steps below.

### 2.1 Assumptions

Here we list the main assumptions in the model and describe the components of the model shown in Figure 1.

- **Geometries:** We investigate how spine geometry (spine volume, shape, neck geometry) and ultrastructure (spine apparatus) can influence synaptic weight change, with the goal of drawing relationships between these different morphological features and synaptic weight (Figure 1e). **Idealized geometries:** Idealized geometries of thin, mushroom, and filopodia-shaped spines were selected from Alimohamadi *et al.* (*45*) and the different geometric parameters are given in Table S1 and S2, Figure 1b.

– **PSD:** For each control geometry, the Postsynaptic Density (PSD) area was set as a fixed proportion of the spine volume.
– **Size variations:** For each spine geometry, we vary the volume of the control geometry to consider the impact of different morphological features.
– **Spine apparatus:** A spine apparatus is included in the thin and mushroom idealized spines by scaling the spine geometry to a smaller size and including it within the plasma membrane geometry. These variations were included to modify the volume of the spine in the presence of these organelles. **Realistic geometries:** We also investigated how spine morphology affected synaptic weight change in realistic morphologies. Realistic spine morphologies were reconstructed from 3D electron microscopy images (*51*) to sufficient mesh quality to import into MCell (*52*). Realistic spines were selected to have a variety of morphologies to reflect filopodia-shaped, thin, and mushroom spines. PSD were denoted based on the segmentation of the original 3D electron micrographs.
- **Timescales:** Our goal is to consider the initial changes in synaptic weight due to a single calcium pulse consistent with prior studies (*10, 37*). Our focus is on early timescale events associated with synaptic weight, rather than the induction of LTP/LTD specifically. The timescale of calcium transients is rapid, on the millisecond timescale (*37, 39, 47*), due to the single activation pulse, various buffering components, and the Spine Apparatus (SpApp) acting as a calcium sink, rather than a source (*53, 54*). Each spine geometry is initiated with a basal concentration of calcium as shown in Table 2. For each geometry, these concentrations were converted to numbers of Ca^2+^ ions to initialize the particle-based simulations.
- **Calcium model stimulus:** The stimulus used in the model is a Excitatory Postsynaptic Potential (EPSP) and Back Propagating Action Potential (BPAP) offset by 10 ms, and a glutamate release that activates *N*-methyl-D-aspartate Receptor (NMDAR) (*37*) as shown in Figure 1a, inset. We include the presynapse as a surface from which glutamate is released from a central location. *α*-amino-3-hydroxy-5-methyl-4-isoxazolepropionic Acid Receptor (AMPARs) which competes with NMDARs to bind glutamate are also included in the model, but do not contribute to the calcium influx.
- **Channel and receptor dynamics:** We assume that the surface density of the receptors and channels on the membrane of the spine is constant and uniformly distributed (*28, 55*). This assumption is based on experimental observations (*55*) and has been employed in other computational models of calcium dynamics (*35, 56, 57*). An important consequence of this assumption is that when the surface area of the spine changes, the total number of receptors will also change. How calcium influx scales with spine volume is an important consideration with implications on how calcium concentration scales with spine size (*12*). The constant receptor density assumption means that calcium influx under-compensates for increases in spine volume (*12*).
- **Boundary conditions:** Calcium ion influx occurs through NMDAR localized to the PSD region and Voltage Sensitive Calcium Channels (VSCCs) on the plasma membrane, based on (*37*). Calcium ions leave the spine volume through the pumps on the plasma membrane, Plasma Membrane Ca^2+^-ATPase (PMCA) and Sodium-Calcium Exchanger (NCX), and into the SpApp when present through Sarco/Endoplasmic Reticulum Ca^2+^-ATPase (SERCA) (*35*). We consider the spine as an isolated geometric compartment and do not consider the effect of calcium influx from the dendrite at this timescale. The base of the spine neck has a Dirichlet boundary condition of calcium clamped to zero and acts as a calcium sink, which represents Ca^2+^ leaving the spine into the dendrite due to the sudden increase in calcium in the spine (*47*).
- **Buffers:** We do not model the different buffer species but rather use a lumped parameter approach as was done before (*18, 35*). Since free Ca^2+^ is rapidly buffered in cells, we consider both mobile buffers in the cytoplasm and immobile buffers on the plasma membrane (*35, 58, 59*). We also include an exponential decay of calcium throughout the cytoplasm to capture the complex cytosolic buffering dynamics without including explicit buffers. Addition of more species introduces many more free parameters and can make the model computationally intractable; therefore we focus on a lumped parameter approach.
- **Stochastic trials:** Each simulation condition was run with 50 random seeds and these individual runs were averaged to obtain mean and standard deviation (*37, 46*).
- **Model readouts:** We report Ca^2+^ dynamics in terms of the number of ions rather than concentration. This is because the total number of ions in the spine reflects total signal coming into the spine and is the natural output from these particle-based simulations. The total number of calcium ions is used as input to calculate the synaptic weight change.
- **Synaptic weight:** The synaptic plasticity model developed by Shouval *et al.* (*10*) was adapted to have dependence on total calcium ions rather than calcium concentration. The rate of synaptic weight update depends on a learning rate, *τ_w_*, and a thresholding function, Ω*_w_*, that are both dependent on calcium ion levels, Figure 1c-d. The learning rate determines the rate of synaptic weight change, while Ω*_w_* determines if the weight increases or decreases. Thresholds for Long Term Potentiation (LTP) and Long Term Depression (LTD), *θ_P_* and *θ_D_*, are set so an intermediate level of calcium leads to a weakening of a synapse and LTD, while an elevated level of calcium leads to the strengthening of a synapse and LTP (*10, 60, 61*).

**Figure 1:**
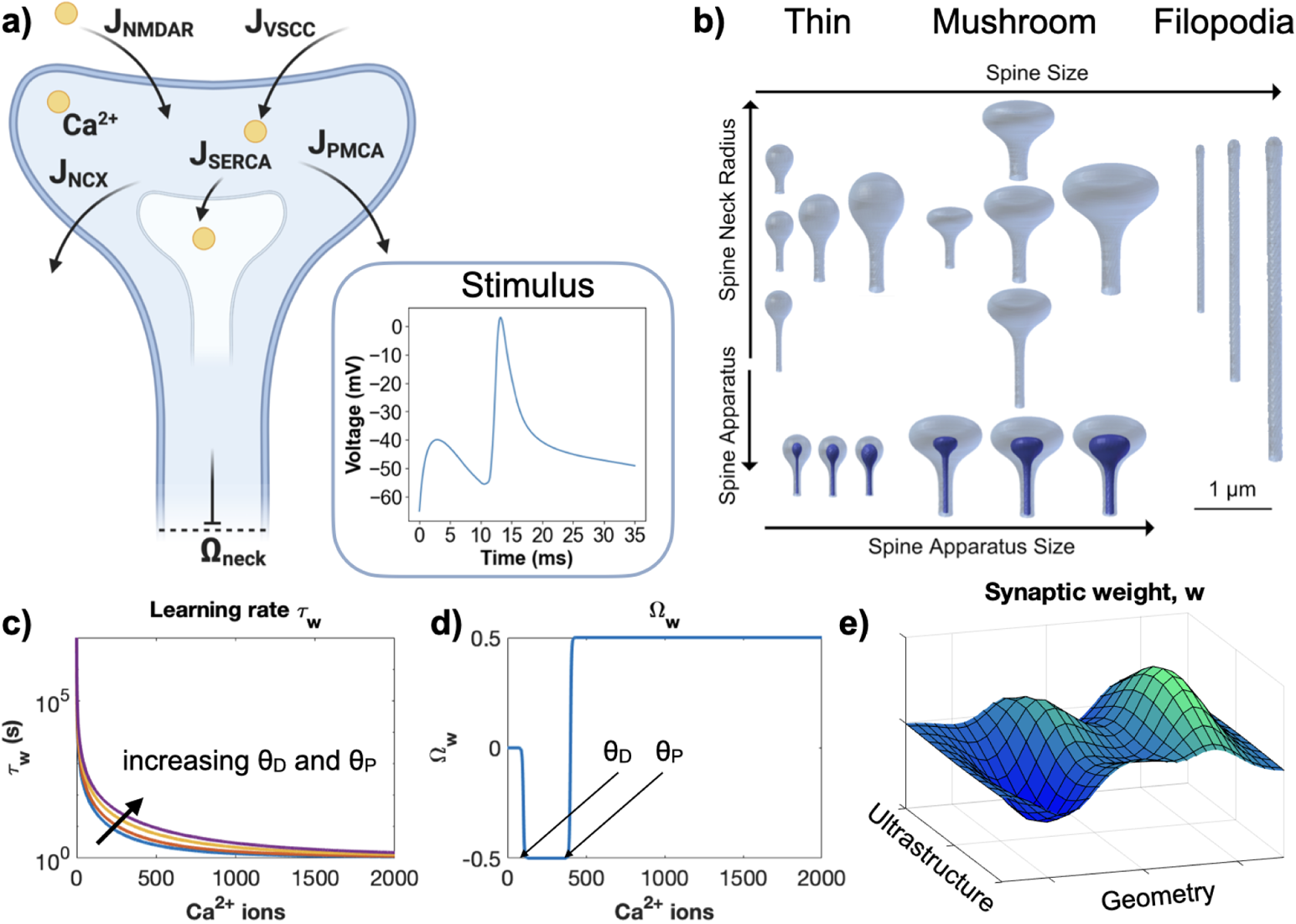
Model overview. (a) Our spatial particle-based model includes calcium influx through NMDAR and VSCC, calcium efflux to the extracellular space through PMCA and NCX pumps, and to the Spine Apparatus through SERCA pumps. Arrows indicate the movement of Ca^2+^ through the labeled pump, channel, or receptor. Ω*_neck_* represents the Dirichlet boundary condition at the base of the spine neck, at which the concentration of calcium ions is clamped to zero. Cytosolic calcium is buffered using cytosolic mobile and membrane-bound immobile calcium buffers. Inset: A change in membrane potential triggered by an excitatory postsynaptic action potential (EPSP) and back propagating action potential (BPAP) acts as the model stimulus, along with the release of glutamate molecules. b) The geometric factors considered in our model include spine shape, spine size, neck radius and length, and spine apparatus (SpApp) size. We investigate three spine shapes: thin, mushroom, and filopodia-shaped. Calcium levels determine the learning rate *τ_w_* (c), and function Ω*_w_* (d), that in turn determine synaptic weight (e). The influence of geometry (spine volume, surface area, PSD area, etc.) and ultrastructure (spine apparatus, internal organelles, etc.) on calcium signaling thus has an influence on synaptic weight. *θ_D_* and *θ_P_* represent the thresholds for long term depression and potentiation, respectively. Panel a) was generated using biorender.com.

**Table 2:**
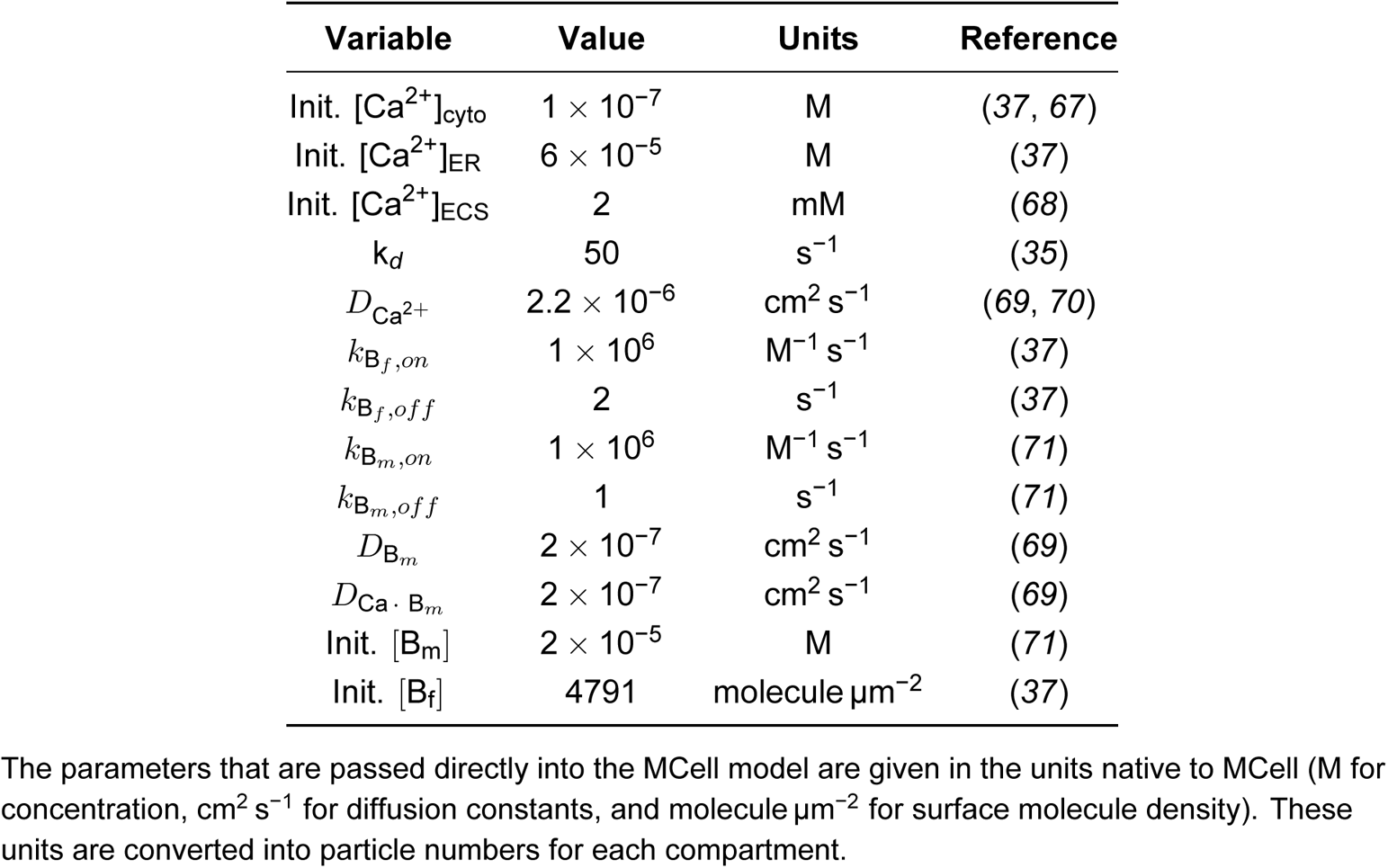
Parameters used in the model.

| Variable | Value | Units | Reference |
| --- | --- | --- | --- |
| Init. $[\text{Ca}^{2+}]_{\text{cyto}}$ | $1 \times 10^{-7}$ | M | (37, 67) |
| Init. $[\text{Ca}^{2+}]_{\text{ER}}$ | $6 \times 10^{-5}$ | M | (37) |
| Init. $[\text{Ca}^{2+}]_{\text{ECS}}$ | 2 | mM | (68) |
| $k_d$ | 50 | $\text{s}^{-1}$ | (35) |
| $D_{\text{Ca}^{2+}}$ | $2.2 \times 10^{-6}$ | $\text{cm}^2 \text{ s}^{-1}$ | (69, 70) |
| $k_{\text{B}_f, \text{on}}$ | $1 \times 10^6$ | $\text{M}^{-1} \text{ s}^{-1}$ | (37) |
| $k_{\text{B}_f, \text{off}}$ | 2 | $\text{s}^{-1}$ | (37) |
| $k_{\text{B}_m, \text{on}}$ | $1 \times 10^6$ | $\text{M}^{-1} \text{ s}^{-1}$ | (71) |
| $k_{\text{B}_m, \text{off}}$ | 1 | $\text{s}^{-1}$ | (71) |
| $D_{\text{B}_m}$ | $2 \times 10^{-7}$ | $\text{cm}^2 \text{ s}^{-1}$ | (69) |
| $D_{\text{Ca} \cdot \text{B}_m}$ | $2 \times 10^{-7}$ | $\text{cm}^2 \text{ s}^{-1}$ | (69) |
| Init. $[\text{B}_m]$ | $2 \times 10^{-5}$ | M | (71) |
| Init. $[\text{B}_f]$ | 4791 | molecule $\mu\text{m}^{-2}$ | (37) |

The parameters that are passed directly into the MCell model are given in the units native to MCell (M for concentration, $\text{cm}^2 \text{ s}^{-1}$ for diffusion constants, and molecule $\mu\text{m}^{-2}$ for surface molecule density). These units are converted into particle numbers for each compartment.

### 2.2 Dynamics of calcium ions in the spine volume

We summarize the main reactions for Ca^2+^ in the volume. These reaction models were obtained from Bartol *et al.* (*37*) and Bell *et al.* (*35*) and are discussed in detail below. Model parameters are given in Table 2. We find that our calcium dynamics are comparable to previously published models (*35, 37, 62, 63*) and experimental observations (*64–66*), Figure S8.

In the volume, calcium binds with fixed and mobile buffers in the cytoplasm, modeled here generically by B*_m_* to represent mobile calcium buffers, and B*_f_* representing fixed buffers. Calcium-buffer binding is modeled by,

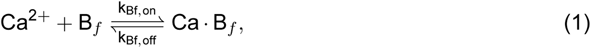

and

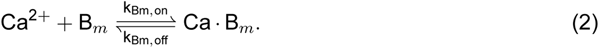

Reaction rates for mobile and fixed buffers are provided in Table 2.

An additional Ca^2+^ decay term is given by

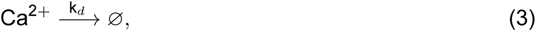

where k*_d_* sets the decay rate. The value of k*_d_* is taken as 50 s^−1^ inspired by (*35, 36*).

### 2.3 Plasma Membrane

The primary influx of calcium through the plasma membrane occurs through NMDARs and VSCCs, and calcium is pumped out of the volume through PMCA and NCX. In this model, NMDARs depend both on voltage and glutamate and are localized to the PSD region. VSCCs are voltage-dependent and localized throughout the plasma membrane. PMCA and NCX are calcium-dependent pumps and are also located throughout the plasma membrane surface.

#### 2.3.1 NMDA receptors

NMDARs are localized to the PSD area with a surface density of 150 molecule µm^−2^ (*37*). The activation of NMDAR is modeled with an asymmetric trapping block kinetic scheme as proposed in Ref. (*72*). The activation of NMDAR depends on the diffusion of glutamate through the synaptic cleft and its binding to inactive receptors. A surface identical to the top of the spine head is translated 2 µm above the head to represent the synaptic cleft. At time t = 0 in each simulation, 500 molecules of glutamate are released at the center of this synaptic cleft. The released glutamate molecules diffuse through the cleft at a rate of 2.2 × 10^−6^ cm^2^ s^−1^ and bind to membrane-bound proteins. On the postsynaptic membrane, NMDARs compete with the glutamate receptor AMPAR for glutamate; thus, AMPARs are also included in the simulation to model this competition, but do not play a role in calcium influx. AMPAR is also localized to the PSD area at a density of 1200 molecule µm^−2^ (*37*). The binding of glutamate to AMPAR is modeled according to the kinetic scheme proposed in Ref. (*73*).

Calcium ion flux through open NMDARs is modeled by the simple unimolecular reaction,

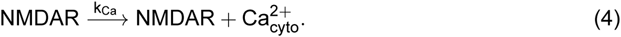

The rate of calcium influx is given by

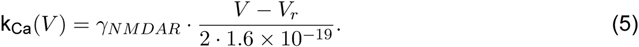

Here, V is the membrane potential, and V_r_ is the reversal potential of NMDAR. The parameters for the NMDAR reactions are the same as given in (*72*) and the parameters for the AMPAR reactions are the same as those in (*73*).

#### 2.3.2 Calcium influx through voltage-sensitive calcium channels

The influx of Ca^2+^ through an open VSCC is given by the reaction:

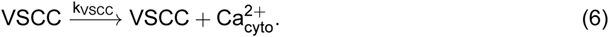

The rate of calcium influx is given by

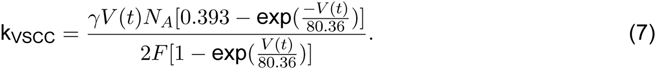

The influx of Ca^2+^ through VSCCs is also dependent on the activation kinetics of VSCCs. The initial conditions for all the VSCCs is the closed state, and the activation of the channels is modeled here with a five state kinetic scheme as used in Ref. (*37*). The parameters for Ca^2+^ influx through VSCCs are the same as in Ref. (*37*). VSCCs are located on the Plasma Membrane (PM) with a density of 2 molecule µm^−2^.

#### 2.3.3 Voltage calculations in the model

Since the transmembrane potential varies with time (Figure 1a, inset) and the rate constants for NMDAR and VSCC are voltage-dependent, the values of these rate constants at each simulation step were precomputed and passed into MCell. The voltage stimulus representing a single EPSP starting at time t = 0, followed by a single BPAP occurring at an offset of 10 ms was obtained from Ref. (*37*). Note that this time offset is within the typical window for Spike-Timing Dependent Plasticity (STDP) to induce LTP (*37, 69*).

#### 2.3.4 PMCA and NCX dynamics

PMCA and NCX are located on the plasma membrane with areal density 998 molecule µm^−2^ and 142 molecule µm^−2^ respectively (*37*), forcing calcium to flow out of the cell. These pumps are modeled using the set of elementary reactions and reaction rates from Ref. (*37*).

#### 2.3.5 Spine Apparatus

Calcium enters the spine apparatus via SERCA pumps, and leaks out. SERCA pumps are calcium-dependent and located throughout the spine apparatus membrane at 1000 molecule µm^−2^ (*37*). SERCA influx is modeled as a series of elementary reactions with rates from Ref. (*37*). Calcium leakage from the spine apparatus into the cytosol is modeled by the reaction

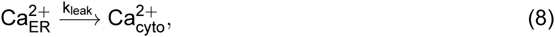

where k_leak_ is 0.1608 s^−1^ from Ref. (*35*).

### 2.4 Extracellular calcium

Extracellular calcium was not explicitly modeled for ease of computational tractability. We assumed a constant extracellular calcium concentration (2 mM) that is negligibly impacted by the calcium influx to and efflux from the spine cytoplasm. The dynamics of Ca^2+^ are explicitly modeled once they enter the cell through channels located on the PM, and cease to be explicitly represented once they are pumped out of the cell.

### 2.5 Synaptic weight change

Synaptic weight update was calculated using the classical model from Shouval *et al.* (*10*). The governing equations were modified to take total number of Ca^2+^ ions rather than a concentration. Here we use total number of ions to highlight the details available from a stochastic simulation and to consider the consequences of using this global readout on synaptic weight.

We modeled the changes in synaptic weight, *w*, as a phenomenological relationship, inspired by (*10, 18*). In our model, the change in synaptic weight is given by

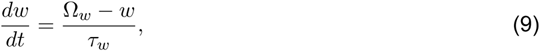

where *τ_w_* is a learning rate given as

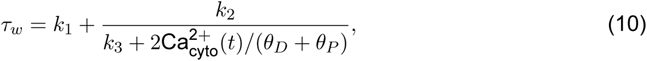

and Ω*_w_* describes calcium dependence in the regimes of LTP and LTD as

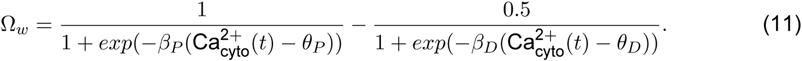

We note that cytosolic calcium, 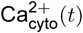, is the total number of ions in the spine. The differential equation for synaptic weight, *w*, is solved in MATLAB 2018b using ode23s, for each calcium transient predicted by the MCell model. The initial synaptic weight value is set to 0 so the change in synaptic weight and synaptic weight are the same value for this single stimulation event. Synaptic weight parameters are given in Table 3.

**Table 3:**
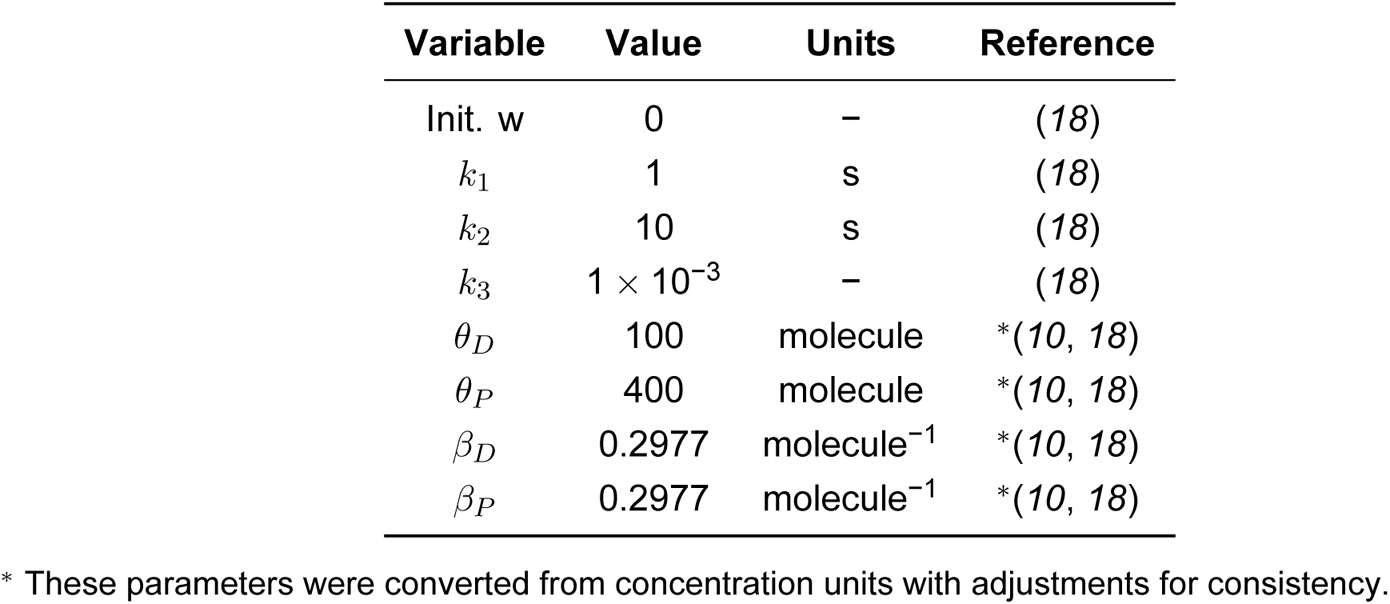
Parameters for Synaptic Weight.

### 2.6 Simulation Information and Parameters

Calcium simulations were conducted for a total simulation time of 35 ms with a 500 ns time step. Each geometry is simulated in MCell with 50 distinct seeds to generate an appropriate sample size of results. All simulations use a write-out frequency of once per iteration for reproducibility of results. Longer write out frequencies introduce non-determinism to the trajectories arising from the MCell reaction scheduler. At the beginning of each simulation, membrane proteins are randomly distributed over specified regions of the spine geometry surface area according to an assigned count or concentration.

System configuration and analysis scripts are all available on Github https://github.com/RangamaniLabUCSD/StochasticSpineSimulations.

## 3 Results

We first demonstrate the coupling between stochastic dynamics of calcium in a single spine to the deterministic model of synaptic weight update (Figure 2). We then investigate whether spine size has any effect on synaptic weight change of filopodia-shaped spines (Figure 3), thin spines (Figure 4), and mushroom spines (Figure 5). Next, we consider the role of the spine apparatus (Figure 6). Finally, we investigate the relationship between spine morphology and synaptic weight update in realistic geometries (Figure 7). Our results predict that synaptic weight change through calcium dynamics is a deterministic function of geometric parameters of the spines (Figure 8). We discuss these results in detail below.

**Figure 2:**
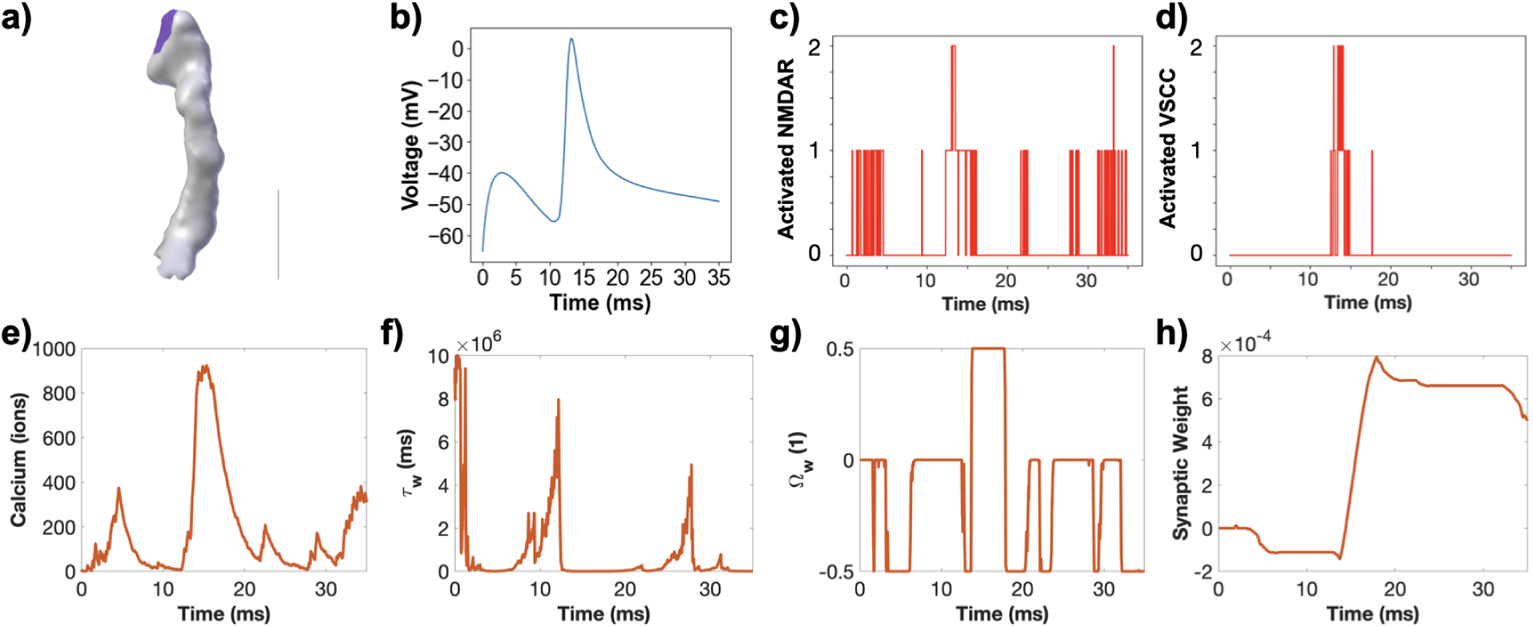
Stochastic receptor, channel, and calcium dynamics inform deterministic synaptic weight update. a) A realistic thin spine with a volume of 0.045 µm^3^ serves as an example spine to consider how stochastic receptor and channel dynamics translate into calcium transients that inform synaptic weight update. Scale bar: 0.5 µm. b) The model stimulus includes a set voltage profile that activates both NMDARs and VSCCs. We consider a single seed run (seed 1 for the realistic thin spine 39). c) Activated, open NMDARs over time for a single simulation in the realistic geometry shown in (a). d) Activated, open VSCCs over time for a single simulation in the realistic geometry shown in (a). e) Calcium transient due to the channel and receptor dynamics shown in (c-d). Learning rate *τ_w_* (f), Ω*_w_* (g), and synaptic weight update (h) are calculated from the calcium transient in (e).

### 3.1 The coupled model filters stochastic Ca^2+^ dynamics to synaptic weight update

We demonstrate how the stochastic calcium model informs the deterministic synaptic weight predictions. We consider a single seed trial for a single geometry, in this case seed 1 from a realistic thin spine (Figure 2a). We observe that in response to the voltage trace and glutamate release (Figure 2b), the NMDARs and VSCCs stochastically open and close (Figure 2c, d), resulting in a noisy calcium transient (Figure 2e). We compare our calcium transients to both published experimental (*64–66*) and computational (*35, 37, 62, 63*) results and find a reasonable agreement, Figure S8. We next calculate the learning rate (Figure 2f) and Ω*_w_* (Figure 2g) terms in the synaptic weight model as a function of this calcium transient. We notice that the noisy calcium dynamics are filtered into a smoother synaptic weight prediction (Figure 2h). For another example of how calcium pulse magnitude and width translates to synaptic weight update, see Figure S2. With this understanding of how the two models integrate, we next investigate how spine geometry influences calcium transients and subsequent synaptic weight predictions.

### 3.2 Synaptic weight change depends on spine volume-to-surface ratio in filopodia-shaped spines

We begin our analysis with a simple question – does spine size alter synaptic weight change? To answer this question, we first examined filopodia-shaped spines. Dendritic filopodia are precursors of dendritic spines and serve to bridge the gap between the dendrite and an axon that is passing by during synapse formation (*74*). These are highly motile elongated structures that resemble tubules (lengths of 2–20 µm and neck diameters smaller than 0.3 µm). The simplicity of this geometry allows us to focus on the role of size alone in a simple spine geometry. We used spine geometries of three different volumes (0.017, 0.058 and 0.138 µm^3^). Simulations revealed that the calcium dynamics in these tubule-shaped spines appeared to follow a ‘plug-flow’ behavior where at 15 ms, all the calcium is localized to one region (Figure 3a). This behavior is because of the narrow geometry of the spine, preventing dispersion of the calcium (see also Supplemental Movie S1). Next, we examined the temporal dynamics of calcium and note that the larger spines have larger numbers of calcium ions (Figure 3b) but also have a larger variance of calcium ions (Figure 3c). We further characterized the dynamics by considering the peak calcium values and decay time constants of the calcium transients versus the spine volume-to-surface area ratio. We chose the volume-to-surface area ratio as a geometric metric of spine morphology because it encompasses both the cytosolic volume through which calcium diffuses and the surface area of the spine membrane through which calcium can enter and leave the system. Additional analyses with respect to spine volume are shown in Figure S3.

**Figure 3:**
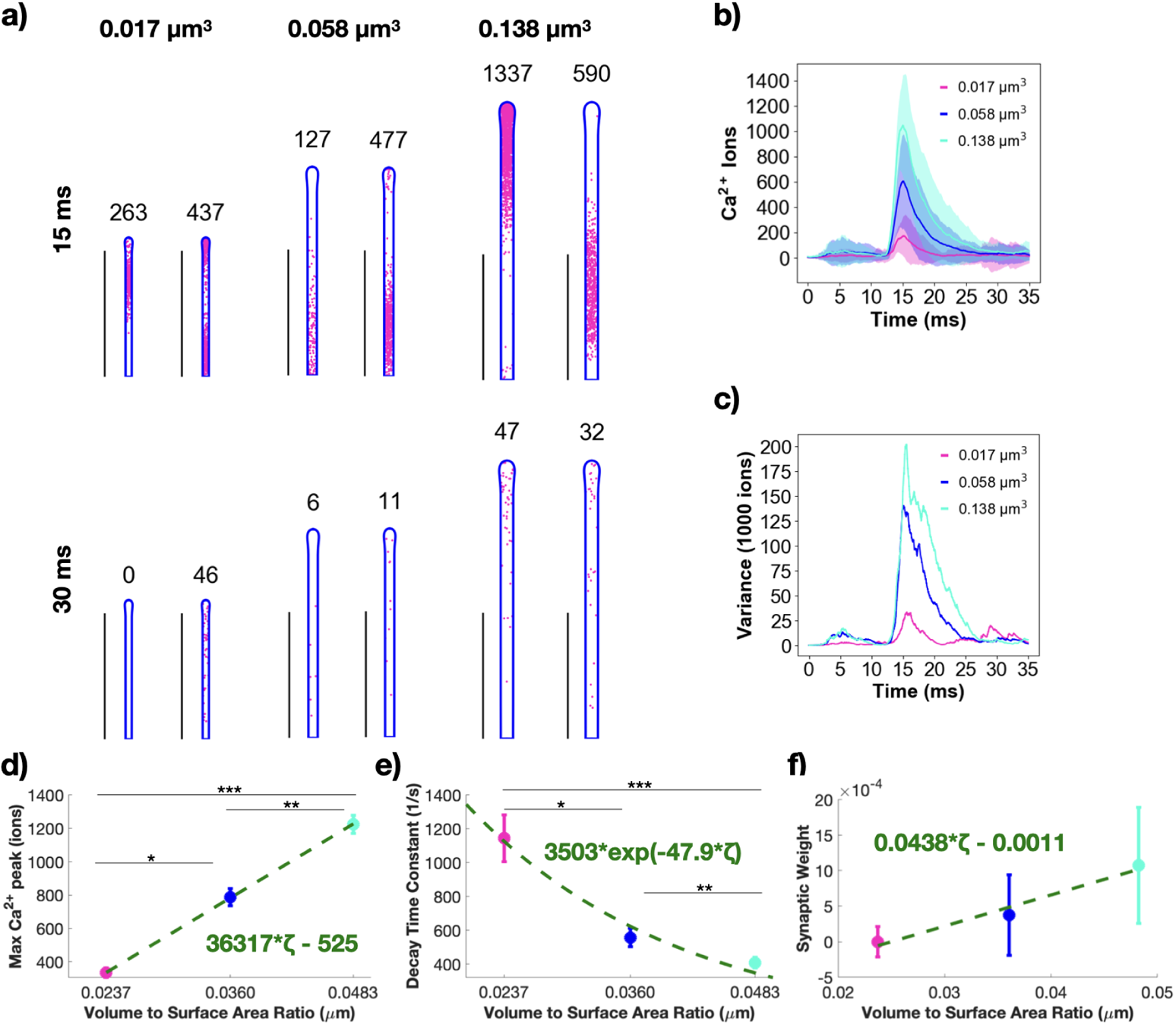
Calcium dynamics and synaptic weight change in filopodia-shaped spines depend on spine volume-to-surface ratio. a) Spatial plots illustrating Ca^2+^ localization at 15 and 30 ms for filopodia-shaped spines with different volumes (0.017, 0.058 and 0.138 µm^3^). The number above each geometry corresponds to the number of Ca^2+^ in that frame. Two random seeds are shown as examples for each geometry. Scale bars: 2 µm. b) Mean (solid) and standard deviation (shaded area) of Ca^2+^ transients across 50 simulations for each of the three filopodia-shaped spine sizes. c) Variance of Ca^2+^ over time, displayed as variance divided by 1000 ions. d) The mean and standard error (n=50) of the peak number of Ca^2+^ in different filopodia-shaped spine sizes shows statistically significant differences; p* = 2.0262 10^−11^; p** = 9.898 10^−8^; p*** = 4.362 10^−26^ using a two-tailed *t*-test. We fit the trend in peak Ca^2+^ as a linear function of volume-to-surface area ratio, *ζ*; *r*^2^ = 0.5521 for the linear fit. e) The decay timescales of each Ca^2+^ transient are estimated by fitting with an exponential decay function *c* exp(−*kt*). The mean and standard error (n=50) of the decay time constant, *k*, shows statistically significant differences across filopodia-shaped spine sizes; p* = 1.6331 10^−4^; p** = 0.0209; p*** = 1.3381 10^−6^ from a two-tailed *t*-test. The mean decay time constants as a function of volume-to-surface area ratio, *ζ*, were fit with an exponential *a* exp(−*bζ*); *r*^2^ = 0.203 for the exponential fit. f) The mean and standard error (n=50) of the calculated synaptic weight change at the last time point in the simulation for all filopodia-shaped spine sizes, plotted against the volume-to-surface area ratio, shows statistically significant differences between all cases; p_12_ = 2.7290 × 10^−5^; p_23_ = 2.8626 × 10^−6^; p_13_ = 1.6321 × 10^−14^ from two-tailed *t*-test, where 1, 2, and 3 refer to the spines in increasing size. We fit the trend in synaptic weight as a linear function of volume-to-surface area ratio, *ζ*; *r*^2^ = 0.3594 for the linear fit.

We note that, indeed, increasing spine size and therefore the volume-to-surface ratio, causes a linearly proportional and significant increase in peak calcium ions (Figure 3d). We also found that the decay time of calcium from the peak decreased with increasing volume-to-surface area ratios and satisfied an exponential dependence (Figure 3e). As spine size increases, the decay time constant decreases, showing that it takes longer for calcium to clear out of the larger spines and spines with larger volume-to-surface area ratios. Finally, we calculated the synaptic weight change (see Supplemental Section 2.5) and compared this value at 35 ms across volume-to-surface area ratios for the filopodia-shaped spines (Figure 3f). We observed that while the smallest spine had no observable weight change presumably because of the net low calcium influx, the weight change increases with increase in spine volume-to-surface-area ratio (Figure 3f). Thus, we find that even for a shape as simple as a filopodia-shaped spine, changes in spine volume-to-surface area ratio can dramatically alter calcium dynamics and synaptic weight change even in stochastic conditions, suggesting a close coupling between spinogenesis and calcium handling.

### 3.3 Thin and mushroom spines modulate synaptic weight changes as a function of volume-to-surface area ratio

We next asked if the relationships of spine size and synaptic weight change observed for filopodia-shaped spines (Figure 3) also hold for thin and mushroom-shaped spines. Thin and mushroom-shaped spines emerge from filopodia-shaped spines as spinogenesis progresses (*74, 75*). While it has been proposed that spines exist in a continuum of shapes (*76*), historically it has been useful to categorize spines into specific categories of shapes (*77*). Thin spines, with small heads and thin necks, have been classified as ‘write-enabled’ or learning spines due to their high motility. Mushroom spines, on the other hand, with bulbous heads and relatively wider necks, are termed ‘write-protected’ or memory spines due to their stability (*78*). Thin spines are characterized by a spherical head and we repeated the calcium influx simulations in a small thin spine with three different spine neck lengths 0.07, 0.06 and 0.04 µm and thin spines of three different volumes (0.035, 0.119 and 0.283 µm^3^) that were informed by the ranges found in the literature, Figure 4. We observe that, in thin spines, the calcium ions are concentrated in the head at 15 ms but disperse more uniformly by 30 ms (Figure 4a-b and Supplemental Movie S2). We do not observe a plug-flow-like behavior as we did for filopodia-shaped spines likely because of the differences in both shape and volume of the thin spines. The thin spines with different neck length showed very similar calcium transients and variance (Figure 4c-d), except for the thin necked spine which showed much more variance during its decay dynamics. For a closer look at the thin spine neck variation dynamics, see Figure S5. Calcium dynamics in thin spines follows the expected temporal dynamics (Figure 4e), with larger spines having larger peak calcium and increased time to decay. Larger thin spines also have larger variance in the calcium ion transients over time (Figure 4f). Next, we found that the maximum calcium ions per spine was significantly larger in larger spines with statistically different values for the different sized spines. The peak calcium increased linearly compared to spine volume-to-surface area but with a smaller slope when compared to the filopodia-shaped spines (max peak values in filopodia-shaped spines increased three times faster than those in thin spines), (Figure 4g). This suggests that the size dependence of calcium grows slower in thin spines than in filopodia-shaped spines. The decay time also showed an exponential decay in thin spines with increasing volume-to-surface area ratio (Figure 4h). The exponent was smaller for thin spines when compared to filopodia-shaped spines (47.9 versus 22.43) suggesting that the decay rate with respect to volume-to-surface area ratio was slower in thin spines. Finally, the synaptic weight change showed an increase with volume-to-surface area ratio in thin spines (Figure 4i) indicating that larger spines are capable of stronger learning outcomes.

**Figure 4:**
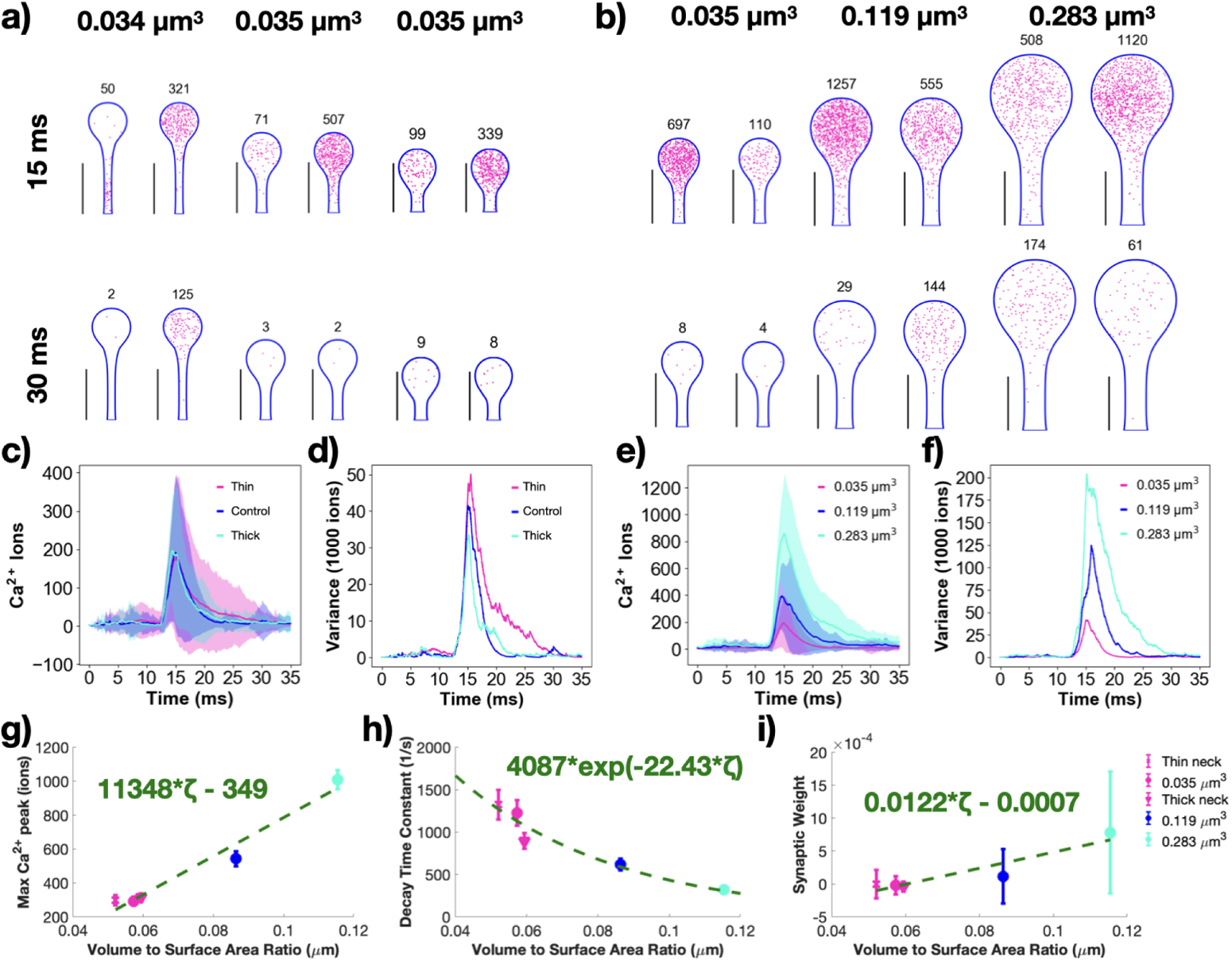
Changing thin spine size modulates calcium dynamics and synaptic weight change. Spatial plots illustrating Ca^2+^ localization at 15 and 30 ms for small thin spines with three different neck lengths 0.07, 0.06 and 0.04 µm (a) and three different volumes 0.035, 0.119 and 0.283 µm^3^ (b). The number above each geometry corresponds to the number of Ca^2+^ ions in the frame. Two random seeds are shown as examples for each geometry. Scale bars: 0.5 µm. c) Mean (solid) and standard deviation (shaded area) of Ca^2+^ transients across 50 simulations for each of the three thin spines with different neck lengths. d) Variance of Ca^2+^ over time for the thin spines with different neck lengths, displayed as variance divided by 1000 ions. e) Mean (solid) and standard deviation (shaded area) of Ca^2+^ transients across 50 simulations for each of the three thin spine sizes. f) Variance of Ca^2+^ over time for the thin spines of different sizes, displayed as variance divided by 1000 ions. g) The mean and standard error (n=50) of the peak number of Ca^2+^ in different thin spine sizes and with different neck lengths show an overall increasing trend. The spines of different sizes show statistically significant differences between the each size; p_12_ = 5.2641 × 10^−6^; p_23_ = 2.7377 × 10^−9^; p_13_ = 5.0036 × 10^−20^ from two-tailed *t*-test, where 1, 2, and 3 denote the different sized spines in increasing size. We fit the trend in peak Ca^2+^ as a linear function of volume-to-surface area ratio, *ζ*; *r*^2^ = 0.4939 for the linear fit. h) The decay timescales of each Ca^2+^ transient are estimated by fitting with an exponential decay function *c* · exp(−*kt*). The mean and standard error (n = 50) of the decay time constant, *k*, shows statistically significant differences across thin spine sizes; p_12_ = 4.3976 × 10^−4^; p_23_ = 1.1541 × 10^−4^; p_13_ = 5.4590 × 10^−8^ from two-tailed *t*-test, where 1, 2, and 3 denote the different sized spines in increasing size. The mean decay time constants as a function of volume-to-surface area ratio, *ζ*, were fit with an exponential *a* · exp(−*bζ*); *r*^2^ = 0.1630 for the exponential fit. i) The mean and standard error (n = 50) of the calculated synaptic weight change at the last time point in the simulation for all thin spine sizes and neck lengths show an increasing trend against the volume-to-surface area ratio. We fit the trend in synaptic weight increase as a linear function of volume-to-surface area ratio, *ζ*; *r*^2^ = 0.2698 for the linear fit. The spines of different sizes shows statistically significant differences between all sizes; p_12_ = 0.0315; p_23_ = 1.0661 × 10^−5^; p_13_ = 2.5751 × 10^−8^ from two-tailed *t*-test, where 1, 2, and 3 denote the different sized spines in increasing size. Inset to right of i: Legend for g-i.

Finally, we repeated our analysis for mushroom-shaped spines of different neck length (0.13, 0.10 and 0.08 µm) and of increasing volume (0.080, 0.271 and 0.643 µm^3^), (Figure 5). The effect of the shape of the spines is evident in the spatial dynamics of calcium (Figure 5a-b and Supplemental Movie S3). Even at 15 ms, we note that while a vast majority of calcium ions are localized in the spine head, there is spillover of calcium into the neck; this is particularly evident in the spines of larger volume in (Figure 5a-b). Spine neck length showed similar increase and decay dynamics to each other (Figure 5c-d), but the thick neck mushroom spine in particular showed a reduced variance. For a closer look at the mushroom spine neck variation dynamics, see Figure S6. The effect of increases in volume, and therefore increases in volume-to-surface area on the temporal dynamics of calcium is an increase in peak calcium (Figure 5e,g) and variance (Figure 5f), and a decrease in the decay time constant (Figure 5h). Interestingly, changing the neck length in the mushroom spine creates the opposite trend with respect to volume to surface area ratio (Figure 5g-h), suggesting that spine neck length is an additional geometric factor for mushroom spines to tune their calcium response. The synaptic weight change in mushroom spines increases with spine volume-to-surface area and is larger for these mushroom spines than the filopodia-shaped and thin spines (Figure 5i). We observe that the peak calcium shows a linear increase with volume-to-surface area ratio with a slope that lies between the thin spines and filopodia-shaped spines. Finally, the decay time constant decreases with spine volume-to-surface area ratio but with a smaller exponential decay when compared to thin spines and filopodia-shaped spines. These two results point to the following conclusions – first, an increase in spine volume results in an increase in critical readouts of synaptic plasticity and second, the shape of the spine alters the quantitative relationships of synaptic plasticity by allowing access to different volume-to-surface area ratios.

**Figure 5:**
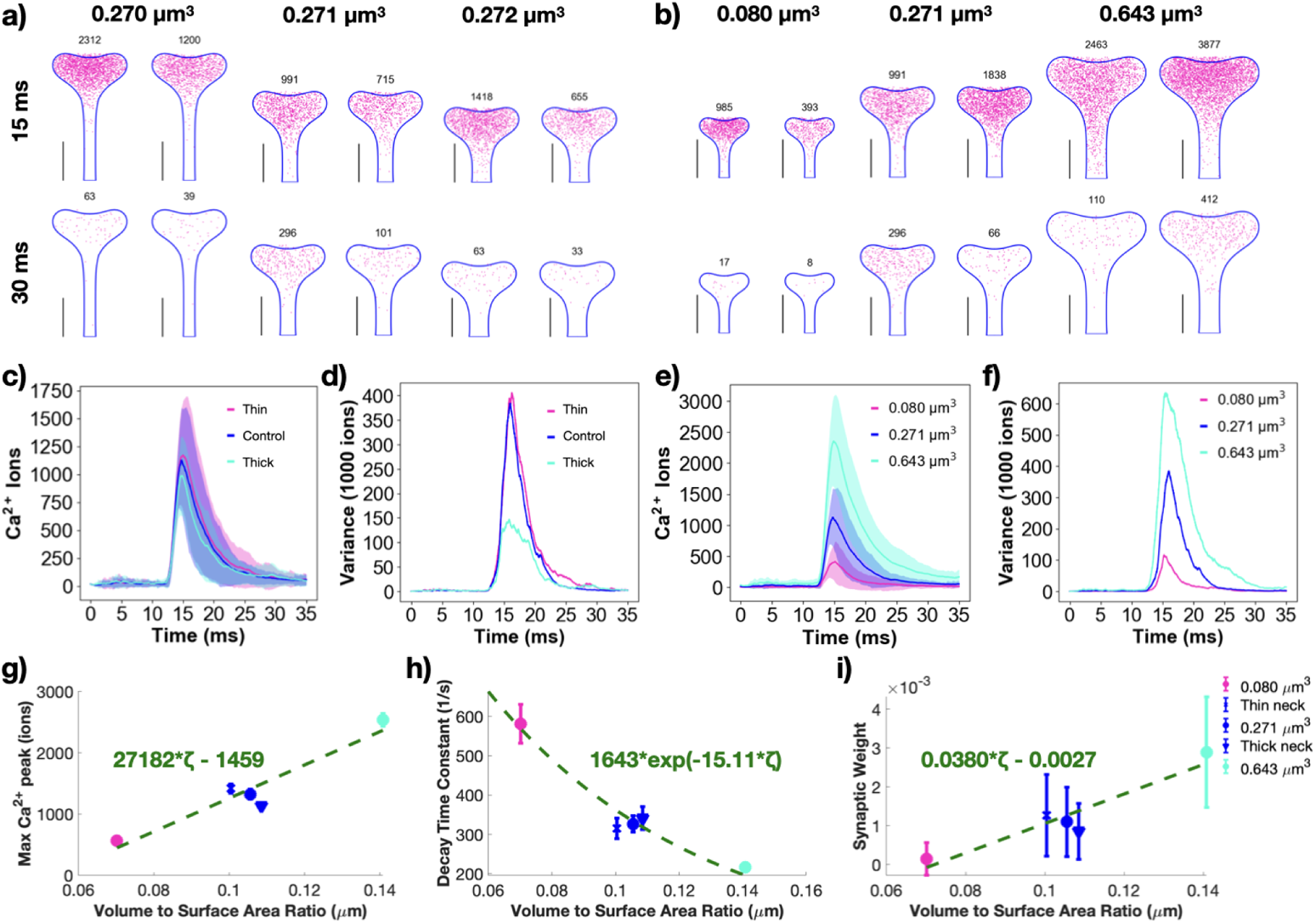
Changing mushroom spine size modulates calcium dynamics and synaptic weight change. Spatial plots illustrating Ca^2+^ localization at 15 and 30 ms for medium mushroom spines with three different neck lengths 0.13, 0.10 and 0.08 µm (a) and three different volumes 0.080, 0.271 and 0.643 µm^3^ (b). The number above each geometry corresponds to the number of Ca^2+^ in the frame. Two random seeds are shown as examples for each geometry. Scale bars: 0.5 µm. c) Mean (solid) and standard deviation (shaded area) of Ca^2+^ transients across 50 simulations for each of the three mushroom spines with different neck lengths. d) Variance of Ca^2+^ over time for the mushroom spines with different neck length, displayed as variance divided by 1000 ions. e) Mean (solid) and standard deviation (shaded area) of Ca^2+^ transients across 50 simulations for each of the three mushroom spine sizes. f) Variance of Ca^2+^ over time for the mushroom spines of different sizes, displayed as variance divided by 1000 ions. g) The mean and standard error (n=50) of the peak number of Ca^2+^ in different mushroom spine sizes and with different neck lengths show an overall increasing trend. The spines of different sizes show statistically significant differences between the each size; p_12_ = 4.1244 × 10^−13^; p_23_ = 6.6467 × 10^−15^; p_13_ = 7.8934 × 10^−32^ from two-tailed *t*-test, where 1, 2, and 3 denote the different sized spines in increasing size. We fit the trend in peak Ca^2+^ as a linear function of volume-to-surface area ratio, *ζ*; *r*^2^ = 0.5474 for the linear fit. h) The decay timescales of each Ca^2+^ transient are estimated by fitting with an exponential decay function *c* · exp(−*kt*). The mean and standard error (n = 50) of the decay time constant, *k*, shows statistically significant differences across mushroom spine sizes; p_12_ = 6.8175 × 10^−6^; p_23_ = 6.4075 × 10^−6^; p_13_ = 1.1118 × 10^−10^ from two-tailed *t*-test, where 1, 2, and 3 denote the different sized spines in increasing size. The mean decay time constants as a function of volume-to-surface area ratio, *ζ*, were fit with an exponential *a* · exp(−*bζ*); *r*^2^ = 0.2380 for the exponential fit. i) The mean and standard error (n = 50) of the calculated synaptic weight change at the last time point in the simulation for all mushroom spine sizes and neck lengths show an increasing trend against the volume-to-surface area ratio. We fit the trend in synaptic weight increase as a linear function of volume-to-surface area ratio, *ζ*; *r*^2^ = 0.4224 for the linear fit. The spines of different sizes shows statistically significant differences between all sizes; p_12_ = 5.1012 × 10^−10^; p_23_ = 2.0097 × 10^−11^; p_13_ = 2.1447 × 10^−23^ from two-tailed *t*-test, where 1, 2, and 3 denote the different sized spines in increasing size. Inset to right of i: Legend for g-i.

### 3.4 Spine apparatus size tunes synaptic weight changes by altering the volume-to-surface area relationships

Approximately 14 % of dendritic spines have specialized endoplasmic reticulum called spine apparatus which are preferentially present in larger, mature spines (*35, 44, 79*). Furthermore, recent studies have shown that the spine apparatus and the ER are dynamic structures in the dendrite and dendritic spines (*80*). Previously, we showed that the spine apparatus modulates calcium transients in deterministic models of calcium influx (*35*) by altering the net fluxes (*36*). Here, we investigate how these relationships are altered in stochastic models in thin and mushroom spines, Figure 6. When a spine apparatus is present in the spine head, it effectively reduces the volume of the spine cytosol and in the time frame of our consideration, acts as a calcium sink (by the action of the SERCA pumps) (*64*). One example trajectory in a mushroom spine with a spine apparatus is shown in Supplemental Movie S10. We varied spine apparatus size in the small-sized thin spine and medium-sized mushroom spine, see Figure 6a-b and Table S2. Calcium transients and variance showed much smoother dynamics for the mushroom spines compared to the thin spines, compare Figure 6e versus c. Peak calcium values were all statistically different for the different spine apparatus sizes in the mushroom spines but not the thin spines. Decay time constants were fit with an exponential relationship (Figure 6h), but there were no statistical differences across different mushroom spines. All different sizes of the spine apparatus produce synaptic weight changes that are statistically different in the mushroom spines; increases in spine apparatus size result in smaller spine volume (and smaller volume-to-surface area ratio) and therefore produce smaller weight changes, Figure 6i. The thin spines had a more complex trend and did not have statistically significant differences in predicted synaptic weight. For a closer look at the variations of the spine apparatus within thin spines, see Figure S7. We conclude that the presence of spine apparatus alters the volume-to-surface area ratio for spines and therefore tunes calcium levels and synaptic weight updates in the large mushroom spines with an inverse relationship to the spine apparatus size.

**Figure 6:**
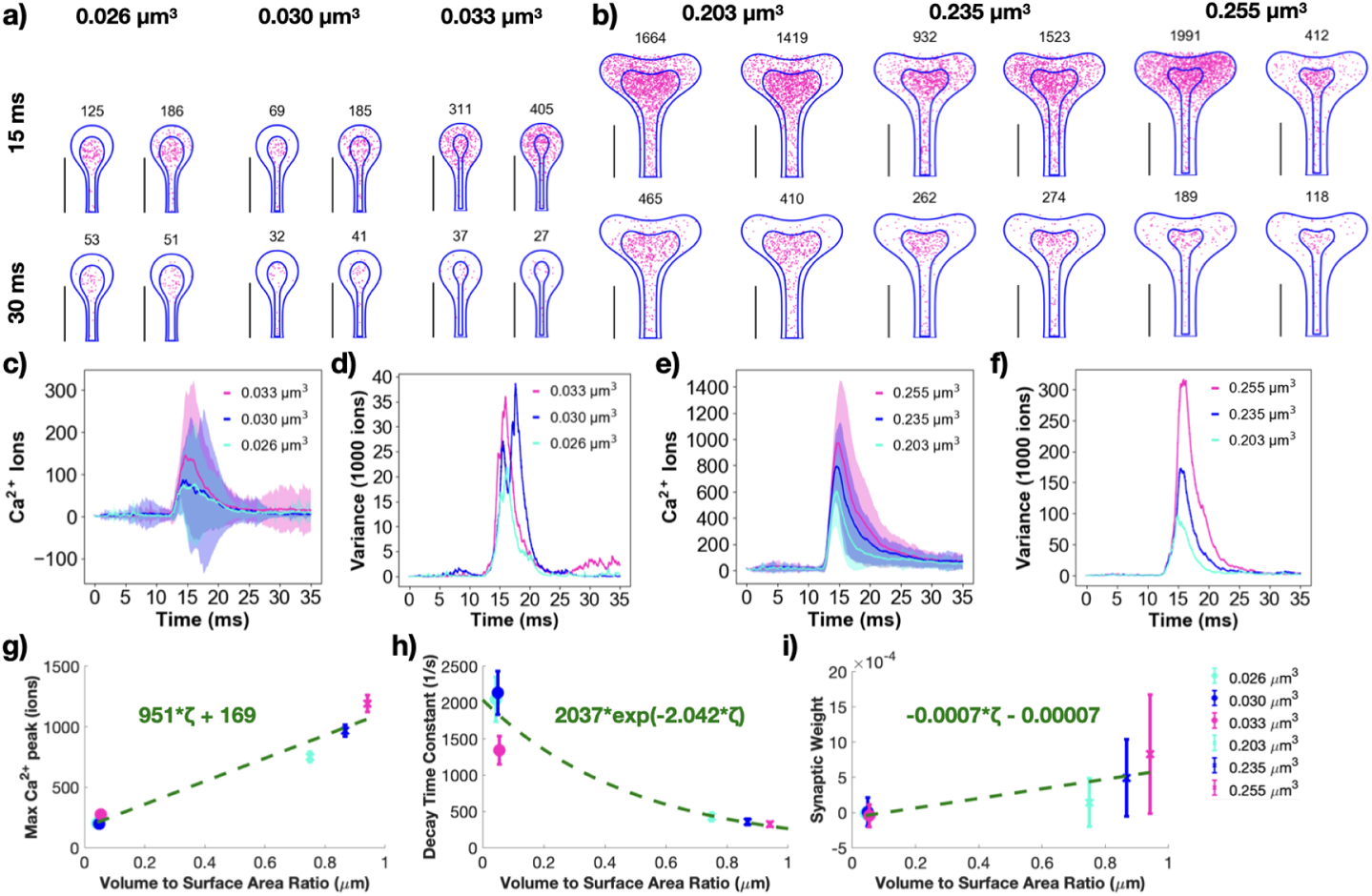
Spine apparatus size modulates synaptic weight change in mushroom spines. a) Spatial plots at 15 and 30 ms for thin spines with spine apparatus of different volumes (net spine volumes of 0.026, 0.030 and 0.033 µm^3^). b) Spatial plots at 15 and 30 ms for mushroom spines with spine apparatus of different volumes (net spine volumes of 0.203, 0.235 and 0.255 µm^3^). The numbers on top of the shape indicate the total number of calcium ions at that instant in both the spine apparatus and cytoplasm. Two random seeds are shown as examples for each geometry. Scale bars: 0.5 µm. Calcium ions over time as mean and standard deviation (c) and variance, displayed as variance divided by 1000 ions (d), for all three thin spines with different spine apparatus sizes. Shaded regions in (c) denote standard deviation. Calcium ions over time as mean and standard deviation (e) and variance, displayed as variance divided by 1000 ions (f), for all three mushroom spines with different spine apparatus sizes. Shaded regions in (e) denote standard deviation. g) Peak calcium ion number for each thin and mushroom spine with a spine apparatus, with the mean and standard error (n=50), show an increasing trend over volume-to-surface area ratio. We fit the trend in peak values with a linear function against the volume-to-surface area ratio, *ζ*; *r*^2^=0.6091 for the linear fit. The mushroom spines show statistically significant differences between its sizes; p_12_ = 0.0010; p_23_ = 0.0101; p_13_ = 4.0801 × 10^−7^ from two-tailed *t*-test, where 1, 2, and 3 denote the different sized spines in increasing cytosolic volume. The thin spines only show statistically significant differences between two of the three paired cases; p_13_ = 0.0453; p_23_ = 0.0461 from two-tailed *t*-test. h) We fit the decay dynamics of each calcium transient with (*c* · exp(−*kt*)) and report the decay time constant, k, as a mean and standard error (n = 50) against volume-to-surface area ratio. The decay time constants were not statistically different for the mushroom spines but the thin spines show statistically difference between the second and third spine; p_23_ = 0.0289 from a two-tailed *t*-test. We fit the trend in decay time constants as a function of volume-to-surface area ratio with an exponential *a* · exp(−*bζ*), where *ζ* is the volume-to-surface area ratio; *r*^2^ = 0.2219 for the fit. i) Calculated synaptic weight change mean and standard error (n = 50) at the last time point for all three thin spines with spine apparatus and all three mushroom spines with spine apparatus show an increasing trend. We fit the trend in synaptic weight with a linear function against the volume-to-surface area ratio, *ζ*; *r*^2^=0.2558 for the linear fit. Calculated synaptic weight change at the last time point for all three thin spines shows no statistically significant difference due to spine apparatus size. The mushroom spines statistically significant differences between all cases; p_12_ = 2.0977 × 10^−4^; p_23_ = 0.0198; p_13_ = 6.0097 × 10^−7^ from two-tailed *t*-test, where 1, 2, and 3 denote the different sized spines in increasing cytosolic volume. Inset to right of i: Legend for g-i.

### 3.5 Simulations in realistic geometries reveals that synaptic weight change depends on spine volume and volume-to-surface area

Thus far, we focused on idealized geometries of spines to identify relationships between key synaptic variables and key geometric variables. We found that the peak calcium value, decay time constant, and synaptic weight depend on the volume-to-surface area ratio within each shape classification. Do these relationships hold for realistic geometries as well? To answer this question, we selected realistic geometries from mesh models (*52*) informed by electron micrographs from Wu *et al.* (*51*).

Realistic spines have more complex geometries that do not fall into the exact morphological categories that we used for idealized spines. To test the significance of these variations, we selected two spines of each shape (thin, mushroom, and filopodia-shaped) and conducted simulations with the exact same parameters as the idealized simulations (Figure 7a). We chose realistic geometries that were within the range of sizes of the idealized geometries. The PSDs in the realistic spines were annotated during the segmentation process and no modifications were made to the PSD marked regions. To capture filopodia-shaped protrusions, we selected long, thin spines (with minimal differentiation between the head and neck) that had marked PSD, because we did not include dendritic filopodia in the dendrite section. Details on how to use realistic geometries in these simulation modalities can be found in the Supplemental Material. We showed the spatial distribution of calcium ions for a single seed for a filopodia-shaped, thin, and mushroom spines (Figure 7b), and found that due to the complexity of realistic morphologies, the calcium distribution was more complicated as compared with those observed in the idealized spines.

**Figure 7:**
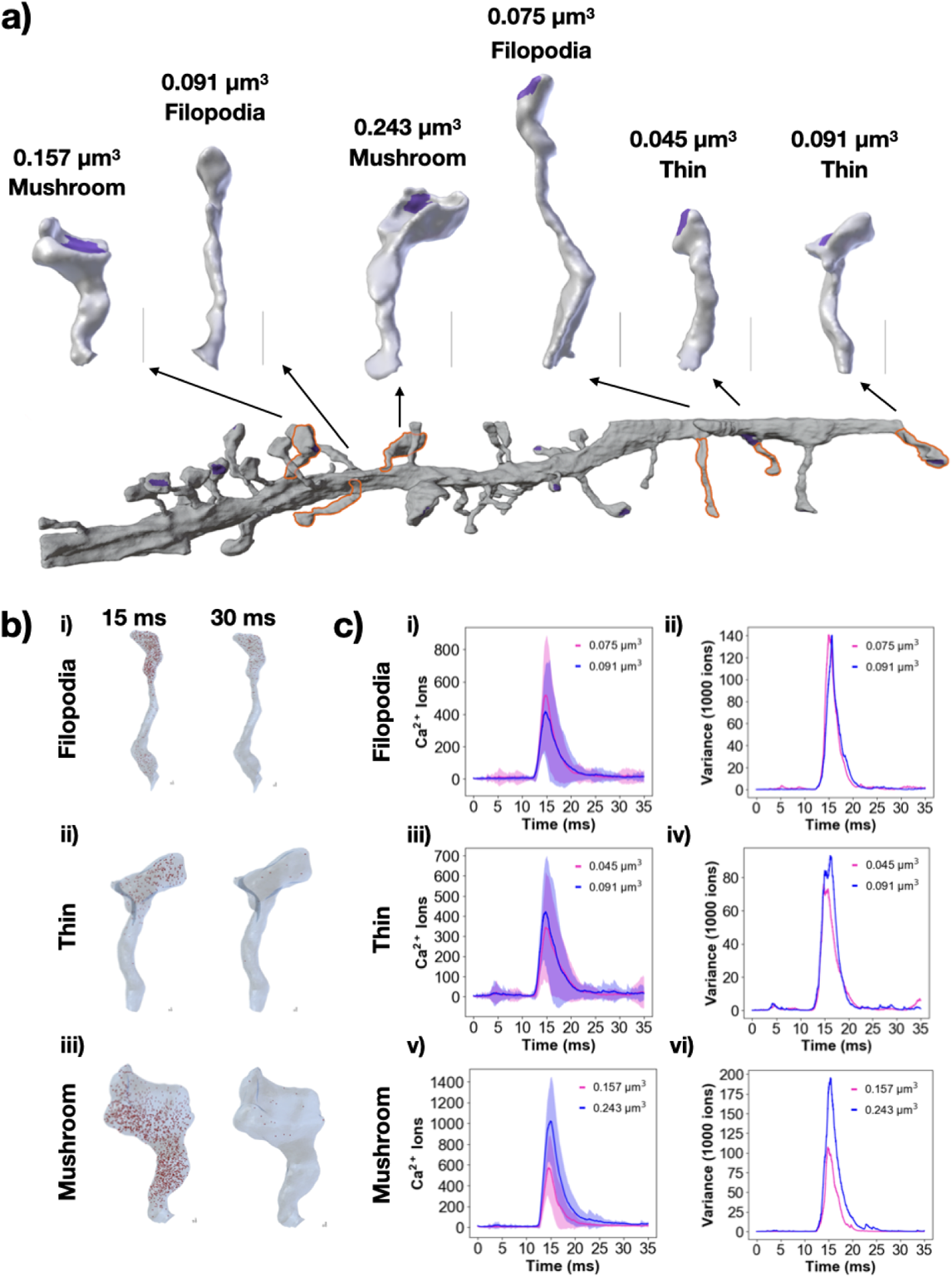
Real spine geometries show size dependence for calcium dynamics. a) Spines similar to the idealized geometries were selected from a reconstructed dendrite (*51*). Representative filopodia-shaped spines, thin spines, and mushroom spines were selected and labelled with their volume and shape. Scale bars: 0.5 µm. b) Snapshots at 15 ms and 30 ms for a single seed for a filopodia-shaped spine (i), thin spine (ii), and mushroom spine (iii). c) Calcium transients as means and standard deviation, along with variance over time for the realistic spines of different shapes; i-ii) filopodia-shaped spines, iii-iv) thin spines, and v-vi) mushroom spines. The realistic spines are labeled with their volumes.

For filopodia-shaped spines, we found that peak calcium and variance varied with volume but the variance was not appreciably different for the two spines that we used to conduct simulations (Figure 7c(i-ii), Supplemental Movie S5, Supplemental Movie S7). The realistic thin spines we chose had volumes similar to the filopodia-shaped spines and they also exhibited calcium dynamics proportional to their volume (Figure 7c(iii-iv), Supplemental Movie S8, Supplemental Movie S9). Mushroom spines had larger volumes and larger PSD areas compared to thin or filopodia-shaped spines (Figure 7c(v, vi), Supplemental Movie S4 and Supplemental Movie S6. Again, the calcium dynamics was proportional to the volume and showed that larger spines have higher peak calcium values. Thus, the relationships of spine geometry and calcium dynamics hold in realistic geometries as well.

## 4 Discussion

Dendritic spines have been extensively studied as biochemical signaling compartments and their role in calcium sequestration has been theorized (*2, 7, 35, 36, 46, 81, 82*). Their unique morphological traits and the classification of spine sizes and shapes with respect to function suggest possible structure-function relationships at the level of individual spines. In this work, we used stochastic modeling of calcium transients in dendritic spines of different geometries to understand how spine size and shape affect the change in synaptic weight. Using a stochastic simulation is important to investigate variance amongst spine shape and size, as dendritic spines have small volumes and probabilistic channel dynamics. Using idealized and realistic geometries, we found that geometric properties, specifically volume-to-surface area, affected key properties of calcium transients including peak calcium, decay time constants, and synaptic weight change. We discuss these findings in the context of different aspects of synaptic plasticity.

Our models predict that despite the individual calcium transients being stochastic, there is a predictive deterministic trend that appears to carry through the different sizes and shapes of the spines used in our model (Figure 8). One of the advantages of our modeling approach here is that we can directly compare across the entire range of idealized and realistic geometries. By considering all the data from our models, for a total of 18 geometries with 50 simulations in each, we find that the peak calcium number is more-or-less linear with the volume, surface area, and volume-to-surface area ratio (Figure 8d,g,j). The decay time constant for calcium transients shows an exponential decay for larger volume-to-surface ratios, volume, and surface area with quite a bit of variability for smaller ratios (Figure 8e,h,k). We note that both peak calcium and decay time constants show clearer trends within the same spine protrusion type (ie. comparing within the same color). And finally, the synaptic weight change increases as volume-to-surface area, volume, and surface area increase (Figure 8f,i,l). We emphasize that our goal is to demonstrate a trend in the data as opposed to building numerical functions. Although we fit the various data, we note that the r^2^ is often weak, indicative of the complexities that underlie such efforts.

**Figure 8:**
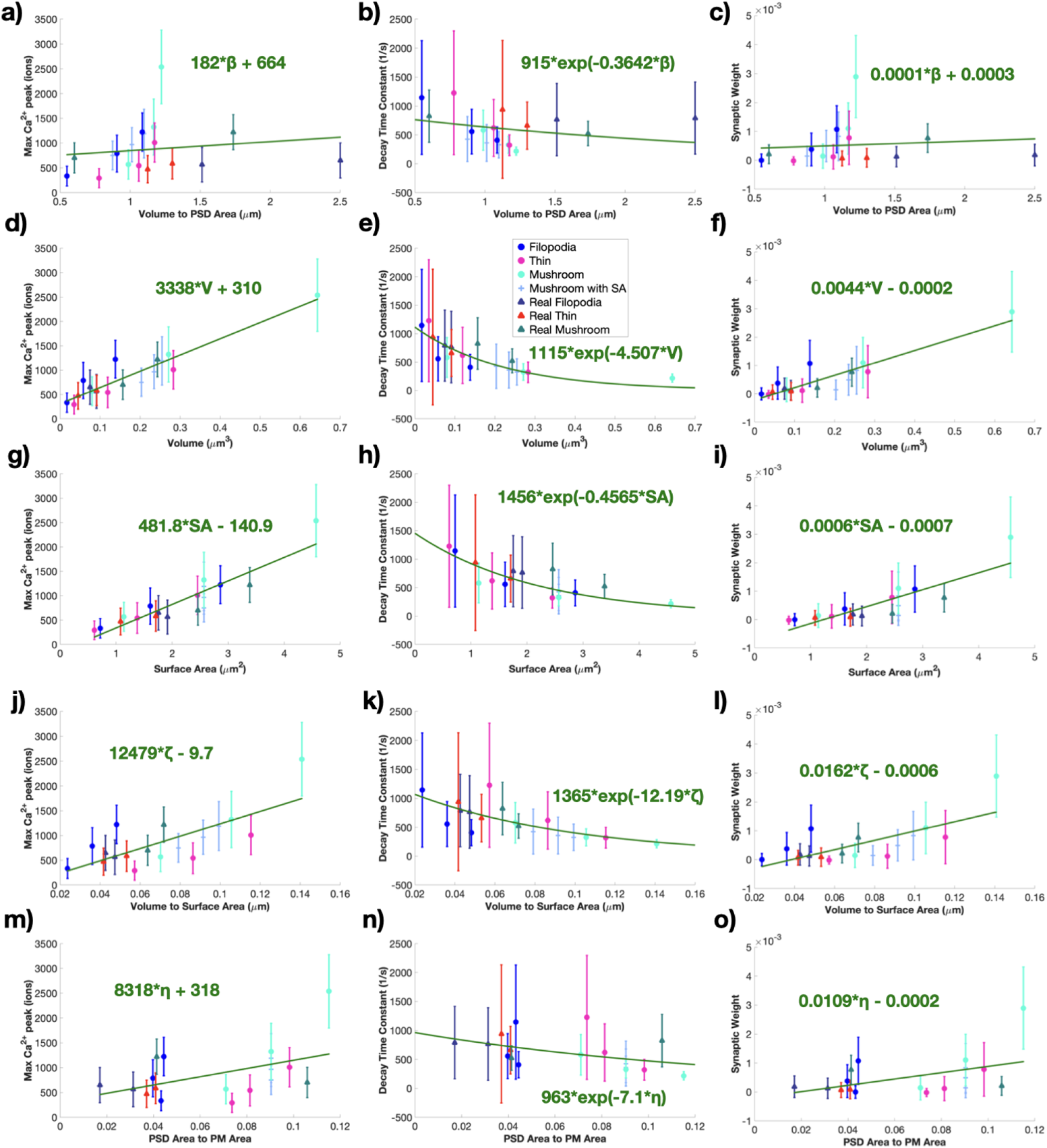
Idealized and realistic spines show overall trends in peak calcium, decay rates, and synaptic weight change with respect to various geometric parameters. We plot all calcium peaks, decay time constants, and synaptic weight predictions for the various geometries against different geometric parameters including volume-to-PSD-Area, volume, surface area, volume-to-surface area ratio, and PSD-Area-to-PM-Area. We fit the trends in peak values with a linear function against the geometric parameter. We fit the decay dynamics of each calcium transient with *c* · exp(−*kt*) and report the decay time constant, k, as a mean and standard error (n = 50) against the geometric parameter. We fit the trend in synaptic weight change with a linear function against the geometric parameter. a) All calcium peaks as mean and standard error (n=50) across volume-to-PSD area ratio show no dependence. *r*^2^ = 0.0152 for the linear fit. b) *r*^2^ = 0.0091 for the fit of decay time constants against volume-to-PSD area ratio. c) Calculated synaptic weight change mean and standard error (n = 50) at the last time point for all idealized and realistic spines shows no dependence on volume-to-PSD area ratio. *r*^2^ = 0.0060 for the linear fit. d) All calcium peaks as mean and standard error (n=50) across volume show a clear increasing trend. *r*^2^ = 0.5666 for the linear fit. e) *r*^2^ = 0.1478 for the fit of decay time constants against volume. f) Calculated synaptic weight change mean and standard error (n = 50) at the last time point for all idealized and realistic spines shows an increasing trend against volume. *r*^2^ = 0.4635 for the linear fit. g) All calcium peaks as mean and standard error (n=50) across surface area show a clear increasing trend. *r*^2^ = 0.5327 for the linear fit. h) *r*^2^ = 0.1427 for the fit of decay time constants against surface area. i) Calculated synaptic weight change mean and standard error (n = 50) at the last time point for all idealized and realistic spines shows an increasing trend against surface area. *r*^2^ = 0.3887 for the linear fit. j) All calcium peaks as mean and standard error (n=50) across volume to surface area ratio show an overall increasing trend. *r*^2^ = 0.351 for the linear fit. k) *r*^2^ = 0.1114 for the fit of decay time constants against volume-to-surface area ratio. l) Calculated synaptic weight change mean and standard error (n = 50) at the last time point for all idealized and realistic spines shows an increasing trend against volume-to-surface area ratio. *r*^2^ = 0.2815 for the linear fit. m) All calcium peaks as mean and standard error (n = 50) across PSD surface area to plasma membrane surface area ratio show an overall increasing trend. *r*^2^ = 0.1441 for the linear fit. n) *r*^2^ = 0.0428 for the fit of decay time constants against PSD-to-surface area ratio. o) Calculated synaptic weight change mean and standard error (n = 50) at the last time point for all idealized and realistic spines shows an increasing trend against PSD-to-surface area ratio. *r*^2^ = 0.1186 for the linear fit. Inset above e: Legend for all plots.

We want to highlight two takeaways from the synaptic weight trends with respect to volume-to-surface-area ratio. The first takeaway is that within a spine shape group (comparing within a specific color in Figure 8), there are clear increasing trends with respect to volume-to-surface area ratio. The second takeaway is that while there are general trends in the data highlighted by the fit lines in Figure 8, there appear to be three regions with slightly different synaptic weight trends at small, intermediate, and large volume-to-surface area ratios. We discuss the possible consequences of these trends in more detail below.

In the idealized geometries, the PSD area is a manually-fixed proportion of the spine volume but realistic geometries do not have this artificial constraint. Therefore, we redid our analysis using volume to PSD area ratio and PSD area-to-surface area ratios (PSD to PM ratio). Interestingly, we did not see a clear trend within the plots against volume to PSD area ratio (Figure 8a-c). In comparison, the PSD area to PM area ratio shows the same relationships overall as volume to surface area ratio (Figure 8m-o) but this time with clustering of data around some ratios. This indicates that the PSD area is an important additional degree of freedom for synaptic weight change that must be considered for interpretation of geometric features and using realistic geometries with boundary markings allows us to investigate this. It is important to note that there is a lot more variability in the smaller volume-to-surface area ratios suggesting the response of smaller spines may be more variable than larger spines. This feature can work as a double-edged sword – it may provide an advantage during the development of spines or be an disadvantage in the case of loss of spines (*40, 83*).

We interpret our predictions in the context of spine shapes. Filopodia are prevalent during early synaptogenesis and can transition into dendritic spines based on synaptic activity (*74*). Additionally, various disease states produce modified dendritic spines that appear more like filopodia (*84*). The lack of significant weight changes for the smallest filopodia-shaped spine indicates that there is a volume threshold at which filopodia receive enough stimulus trigger synaptic weight change and transition towards more stable, mature dendritic spines. Importantly, the early synaptic weight changes emphasize how the increase in spine volume changes the weight outcome from LTD to LTP. This increase in synaptic weight emphasizes how an increase in spine size can push a thin spine to transition into a stable, larger mushroom spine.

The difference in peak calcium level, decay dynamics, and synaptic weight changes as different spine shapes are scanned across different sizes can also provide insight on spine shape transitions during development and maturation. Filopodia-shaped spines have larger increases in peak calcium levels and synaptic weight updates and faster decreases in decay time constants as their volume-to-surface area ratios and volumes increase, compared to both thin and mushroom spines; Figure 3, Figure 4, and Figure 5. This suggests that filopodia can very quickly alter their calcium levels, and therefore are well-suited for initially identifying possible synaptic partners and subsequently directing resources to those filopodia that are good candidates to transition to dendritic spines (*85*). Once filopodia are established, their linear calcium increase with volume might be unsustainable and might lead to the reduced levels of increase for thin spines of comparable volume-to-surface area (and volume). This suggests that larger stimuli might be necessary to push thin spines towards more excitation, perhaps to prevent excessive numbers of thin spines from maturing and leading to resource depletion and excess neural connectivity (*86*). Mushroom spines once again show more of an increase in synaptic weight as they increase in volume-to-surface area ratio (and volume) but at volumes shifted from the filopodia-shaped spines, perhaps highlighting their role as key communication hubs (*86*). The volume shift seen in mushroom spines versus filopodia-shaped spines might serve to limit the number of mature, highly excitable dendritic spines as both a key neuronal network and resource regulation feature. When the spine apparatus acts as a sink, its presence dampens synaptic weight changes in mushroom spines, potentially acting to stabilize the spine from future changes as suggested by others (*18, 78*).

When considering why these trends hold across volume to surface area ratio, it is important to note that Ca^2+^ influx is through receptors and channels with constant densities at the plasma membrane, or in the case of NMDARs localized to the PSD. Therefore, as spines get larger, they have more surface area and more Ca^2+^ influx, which leads to higher numbers of total Ca^2+^ ions. This increase in total ions due to constant receptor and channel densities explains the increasing trend in peak Ca^2+^ number. When considering decay dynamics, Ca^2+^ efflux is due to pumps of constant density on the plasma membrane or on the spine apparatus. Additionally, Ca^2+^ ions decay everywhere in the cytoplasm, bind to mobile buffers in the cytoplasm and fixed buffers on the plasma membrane, and bind to the spine neck base which acts as a sink. Therefore, since many efflux or binding terms are either on the PM through pumps or at the base of the spine neck, larger volume to surface area ratios mean that ions must diffuse further to reach the neck base or PM, explaining why decay time constants seem to decrease with increasing volume to surface area ratio.

While changing geometric features can occur when spines increase and decrease in volume, they can also modify their volume, surface area, and volume-to-surface-area ratio by having a spine apparatus or through changes in the spine neck geometry. We investigated these additional features (Figure 4, Figure 5, and Figure 6; Figure S5, Figure S6, and Figure S7) and found that spine neck and spine apparatus size had volume dependent effects. The smaller thin spine neck length and spine apparatus variations did not show much influence on peak calcium, decay rate, or synaptic weight, while the larger mushroom spine neck length and spine apparatus variations did have some impact on these readouts. Therefore, there are various means in which a spine can modify its synaptic weight response.

There has been substantial debate on deterministic versus stochastic studies for spine signaling (*87–89*). Numerous studies have looked at the importance of stochastic calcium dynamics (*46, 90*), and we agree that the consideration of stochasticity as noise often leads to efforts to average out its effects (*19*). However, comparing our findings here to our previous deterministic results (*35*) shows that geometric factors play a critical role in determining Ca^2+^ dynamics; both approaches show that Ca^2+^ characteristics depend on the volume-to-surface ratio. Additionally, our hybrid approach of stochastic calcium dynamics and deterministic synaptic weight update is becoming increasingly common (*19, 88, 91*). However, care should still be taken in assuming model type as the dynamics of the species, not just particle number, play an important role in the stochasticity of the system (*87*).

We note that our study is only a small piece of the puzzle with respect to synaptic plasticity. There are many open questions remaining. Of particular interest and needing additional exploration is whether one should use total number of calcium ions or calcium concentration in evaluating synaptic weight change. For instance, we find that when calcium results are converted from total ions to average concentration along with the phenomenological synaptic weight equations, we get different trends in synaptic weight update results, Figure S10 and Figure S11. We note that this model of synaptic weight change has been used previously for concentration studies (*10, 18*). We also observe that converting our previous results (*35*) into total ions shows the same trends for max Ca^2+^ peak and decay time constants as this current study, Figure S9. Thus, a simple unit consideration can lead to conflicting results in spatial models and indicates that we need further discussion and investigation on the structure of phenomenological equations for synaptic weight to understand which factors of calcium dynamics matter and to what degree. Additional investigation is also needed in experimental data to relate fluorescence readouts to concentration or molecule numbers. However, we do compare our calcium transients to previously published experimental and computational results and find a reasonable agreement, Figure S8.

Related to these conflicting findings when considering ion total versus concentration, previous studies have considered the assumptions between calcium influx and spine geometry, more specifically the assumption of how calcium influx scales with spine volume (*12*). Here the constant receptor and channel density assumption leads to an undercompensation scenario where calcium influx does not scale with spine volume, leading to lower calcium concentrations for larger spines. See Figure S4 for examples of our model results in terms of calcium concentration. Because the synaptic weight model depends on calcium influx, this means that when using concentration to determine synaptic weight, larger volume spines have less synaptic weight increase. Therefore, the sublinear calcium influx assumption leads to this discrepancy in synaptic weight predictions based on total calcium ions versus ion concentration. See Figure S10 and Figure S11 for examples of this discrepancy. Further research is needed to determine how exactly calcium influx scales with dendritic spine volume *in vivo*, as it is currently unknown which assumption is correct (*12*). Regardless of the relationship between dendritic spine volume and calcium influx, the spine uses various means to modify its calcium transients, including internal organelles such as the spine apparatus acting as either a calcium source or sink (*92*). More research is needed to explore the relationship of geometry-dependent calcium trends and their consequences on phenomenological synaptic weight predictions.

An additional limitation of this study is the usage of traditional p-values for statistical analysis of the data (see Figure S12 for details on h and p-values), since the statistics field has suggested moving away from null-hypothesis significance testing (*93*). We also note that our current focus is on very early events and these models must be extended to longer time scale events to explore the biochemical and geometric interplay for downstream signaling (*16, 94–96*). Associated with these longer timescale events, calcium often occurs in pulse trains due to high frequency stimulation of the dendritic spine (*97, 98*). We compare synaptic weight predictions for a single calcium transient to those due to a pulse train of activation at a single frequency (Figure S13). However, further investigation should be done to more closely consider the role of stimulus magnitude and frequency on synaptic weight update. In addition, it is important to note that this calcium model and dendritic spine geometries are representative of hippocampal pyramidal neurons. Calcium signaling, dendritic spine structure, and synaptic weight induction is neuron-type specific, and other studies, including some MCell simulations, have investigated calcium signaling and synaptic plasticity in other neuron types (*24, 46, 99, 100*). In some neuron types, including Purkinje cells, calcium release from the ER can play a vital role in calcium dynamics and subsequent synaptic plasticity (*24*); thus, care must be taken when considering different neuron types.

In summary, our computational models using idealized and realistic geometries of dendritic spines have identified potential relationships between spine geometry and synaptic weight change that emerge despite the inherent stochasticity of calcium transients. We predict that dendritic spine morphology alters calcium dynamics to achieve their characteristic functions; in particular, so that filopodia can quickly change their synaptic weight, large mushroom spines can solidify their synaptic connections, and intermediate sized spines require more activation to achieve larger synaptic weight changes. Additionally, we predict that within a certain spine shape, increasing volume (and increasing volume-to-surface area ratio), while assuming receptors and channels are also recruited, allows for a larger future increase in synaptic weight, suggesting that the volume change associated with LTP and LTD serves to reinforce the biochemical changes during synaptic plasticity. Therefore, spine morphology tunes synaptic response. The advances in computational modeling and techniques have set the stage for a detailed exploration of biophysical processes in dendritic spines (*53, 94, 101*). Such efforts are critical for identifying emergent properties of systems behavior and also eliminating hypotheses that are physically infeasible (*33, 102*). Models such as this and others can set the stage for investigating longer timescale events in spines including the downstream effectors of calcium (*16, 17, 103, 104*), and actin remodeling for structural plasticity (*105, 106*).

## Abbreviations

AMPAR: *α*-amino-3-hydroxy-5-methyl-4-isoxazolepropionic Acid Receptor
BPAP: Back Propagating Action Potential
EPSP: Excitatory Postsynaptic Potential
LTD: Long Term Depression
LTP: Long Term Potentiation
NCX: Sodium-Calcium Exchanger
NMDAR: *N*-methyl-D-aspartate Receptor
PM: Plasma Membrane
PMCA: Plasma Membrane Ca^2+^-ATPase
PSD: Postsynaptic Density
SERCA: Sarco/Endoplasmic Reticulum Ca^2+^-ATPase
SpApp: Spine Apparatus
STDP: Spike-Timing Dependent Plasticity
VSCC: Voltage Sensitive Calcium Channel

## 5 Acknowledgements

This work was supported by a National Defense Science and Engineering Graduate (NDSEG) Fellowship to M.K.B., a Hartwell Foundation Postdoctoral Fellowship and Kavli Institute of Brain and Mind Innovative Research Grant #2021-1755 to C.T.L., and Air Force Office of Scientific Research FA9550-18-1-0051 to P.R.. MCell development is supported by the NIGMS-funded (P41GM103712) National Center for Multiscale Modeling of Biological Systems (MMBioS). We thank Dr. Lingxia Qiao and Dr. Ali Khalilimeybodi for comments and proofreading.

## Supplemental Material

### S1 Supplemental Methods

We constructed a stochastic model of calcium influx into dendritic spines of different morphologies. In this model, we have calcium influx through various receptors and channels, and calcium efflux through various pumps localized to the plasma membrane or spine apparatus (a specialized form of endoplasmic reticulum). Each simulation condition was run with 50 random seeds. We then take the stochastic calcium temporal dynamics and input them into deterministic equations for synaptic weight update. Using this framework, we can vary spine size and shape to see how geometric factors influence weight updates. We specifically consider calcium dynamics in terms of total calcium ions.

#### S1.1 Readout considerations

We simulate synaptic weight updates for each of the 50 calcium transients and then take the mean and standard deviation of the synaptic weight prediction.

Because we are working with a stochastic model and are considering Ca^2+^ in terms of ions, we converted the parameters in the synaptic weight equations from units involving concentration to units of molecules, based on average spine volumes and realistic numbers of calcium ions in dendritic spines. It is important to note that using total Ca^2+^ ions is a global view of the dendritic spine while concentration can be considered as more of a local measurement. As mentioned, this synaptic weight change is a phenomenological relationship between Ca^2+^ and synaptic weight which captures the concept of synaptic strength change, and it remains unclear if using ions versus concentration is a better approach for predicting this change. We converted our results into average concentrations by dividing the calcium transients by the respective spine volume, converting our synaptic weight parameters into units of concentration, and rerunning our synaptic weight calculations, Figure S10. Interestingly, we see a different trend within the concentration simulations versus total ion simulations. We believe the relationship between spine volume and calcium influx is leading to these differences, and requires further investigation (*12*). We believe that the synaptic weight model might also be parameter dependent, in particular with regards to *β_P_*, *β_D_*, *θ_P_*, and *θ_D_*. This again brings into question the approach of local versus global calcium readouts and how to capture these effects with the parameters in phenomenological models. Further investigation is required to understand the considerations behind these different approaches.

##### S1.1.1 Multiple pulses of calcium

We consider the case in which a spine is activated multiple times to produce a series of calcium pulses, and predict the synaptic weight update from this series of calcium pulses. To do this, we repeated each calcium pulse 10 times with 35 ms between activation events. We note that this is an assumption that has potential consequences on our results. However, for simplicity, we assume the same calcium transient for each pulse train. We input these calcium pulse trains into the synaptic weight equations.

#### S1.2 MATLAB Analysis of Ca^2+^ transients

We used MATLAB version 2018b to analyze the max Ca^2+^ peak and decay time constants for the stochastic Ca^2+^ results. For each realization of the Ca^2+^ transient, we used the max() function to find the peak Ca^2+^ value and corresponding time. We fit the transient after the peak using the fit() function set to ‘exp1’. The parameters from each fit, corresponding to a realization from a random seed, and statistics such as the mean and standard deviation are computed. The standard error of the mean was found by dividing the standard deviation by the square root of the number of individual trials, in this case 50 trials.

#### S1.3 Statistical Analysis

Statistical significance was determined using a two-tailed two-sample *t*-test assuming equal means and variance (ttest2() function) in MATLAB version 2018b with a significance cutoff at p = 0.05. Statistical comparisons were made between the distributions of observables yielded by the 50 simulations of the compared experimental conditions. Trends in the stochastic results data were fit using all 50 seeds for each of the simulations being considered in the fit. The reported trend lines are estimated using the data from all 50 seeds, as opposed to fitting to the means only. Linear fits and exponential fits were computed in MATLAB using the functions fitlm() and fit(), respectively. We highlight that we are using the classical approach of null-hypothesis significance testing, p-values, and statistically significant verbiage, which has been questioned as perilous and over-simplistic (*93*). We have provided the p-values for each result comparison for closer consideration, Figure S12. The linear and exponential trend lines shown have a range of r^2^ values and are used to show general trends. We emphasize however that in some plots we are fitting to either very few data points or a small domain. Therefore, we reiterate that these factors limit the interpretation of the quantitative nature of the fits.

### S2 Geometries

Idealized, axisymmetric geometries are used to represent the structure of dendritic spines in this study. Three general spine shapes are represented – thin, mushroom, and filopodia-shaped – and each shape is further varied in size and, for the thin and mushroom spines, neck radius.

#### S2.1 Geometry generation

The geometries were generated from 2-dimensional ideal spine profiles obtained from Ref. (*45*) consisting of a series of points (r, z) which form the outline of the respective geometry’s rotational cross-section. Using Netgen/NGSolve version 6.2 (*107*), we revolved these profiles about the z-axis to yield a rotationally-symmetric 3-dimensional spine geometry, Figure S1. In all spine geometries, a circular PSD was centered at the top of the spine head. The PSD area was set as a function of spine volume according to the relationship observed in Ref. (*108*).

**Figure S1:**
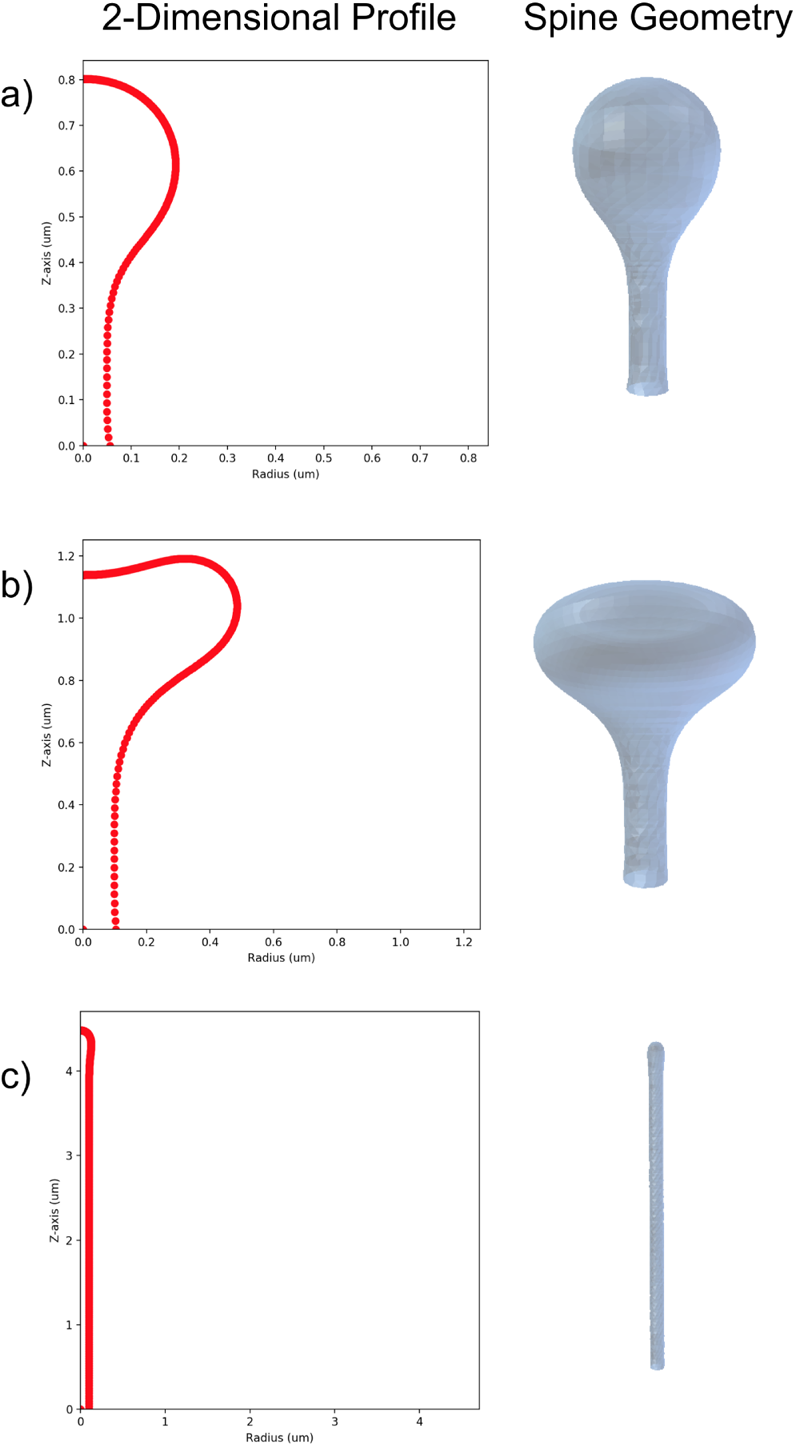
The 2-dimensional spine profiles and the resultant rotationally-symmetric spine geometries for a) thin spines, b) mushroom spines, and c) filopodia.

#### S2.2 Size and neck variations

To further explore the effects of geometric variations on calcium transients and stochasticity, and to facilitate the comparison of spine geometries of similar volumes and different shapes, the base geometries of all three shapes are scaled to two additional volumes beyond the base shapes from (*45*). The additional versions of the thin spine, initially smaller than the other spine shapes, are scaled such that their length measurements are 1.5 and 2 times their original values, resulting in volumes 3.375 and 8 times that of the initial thin spine, respectively. The base mushroom spine, intermediate in volume, is scaled to 0.66 and 1.33 times its original size, resulting in volumes 0.287 and 2.353 times their original value, respectively. And the base filopodia-shaped spine, initially the largest in volume, is scaled to 0.5 and 0.75 times its original size, resulting in volumes 0.125 and 0.422 times the original volume. This scheme ultimately results in three different sizes for each spine shape, spanning a similar range of volumes.

The neck radius of the thin and mushroom spines is also varied, with neck length modified as well to preserve spine volume. To create the different spine sizes, the 2-dimensional spine profiles are dilated about the origin by a certain scale factor, and the resultant image is rotated about its vertical axis using Netgen/NGSolve to produce a scaled-up or scaled-down three-dimensional geometry. In the thin and mushroom 2-dimensional profiles, the x-values of points along the spine neck are scaled by a certain coefficient, and the length of the neck is then scaled by the squared inverse of the coefficient in order to maintain an approximately constant volume. A list of all spine geometries used, and their respective geometric measures, is found in Table S1.

#### S2.3 Spine Apparatus

Some dendritic spines are observed to have a spine apparatus denoted as SpApp, an extension of the smooth endoplasmic reticulum, extending from the dendrite into the neck and head of the spine (*44*). In this study, the effects of the presence of the SpApp on calcium transients and stochasticity are investigated; to achieve this, the thin and mushroom spine geometries are further modified with the addition of a spine apparatus of varying sizes. For both spine shapes, the control-sized SpApp geometry is constructed by scaling down the original spine geometry and extending the spine apparatus neck, such that the SpApp occupies approximately 10 % of the spine volume and extends to the base of the spine. SpApp size is then varied by scaling the SpApp geometry up and down, changing the neck length such that the SpApp base coincides with the spine base. SpApp is not added to the filopodia-shaped geometry, as the spine apparatus is not generally found to be present in such spine shapes (*44*). The SpApp-containing geometries are also listed in Table S2.

#### S2.4 Realistic Geometries

Realistic geometries were chosen from among those on the full dendrite geometry generated in Ref. (*52*). Briefly, the geometric meshes were generated from electron micrographs in Wu et al. (*51*) using GAMer 2 (*109*). Individual spines with labeled PSD and volumes similar to the idealized geometries were selected from the realistic dendritic branch.

### S3 Additional simulation results

#### S3.1 Artificial calcium transients highlight how calcium peak and pulse duration influence synaptic weight prediction

**Table S1:** A list of all geometric variations.

| **Geometry** | **Scale** | **Volume ($\mu\text{m}^3$)** | **Surface Area ($\mu\text{m}^2$)** | **Neck Radius ($\mu\text{m}$)** | **PSD Area ($\mu\text{m}^2$)** |
| --- | --- | --- | --- | --- | --- |
| **Thin** |  |  |  |  |  |
| small | x1 | 0.035 | 0.611 | 0.06 | 0.045 |
| thin neck | x1 | 0.034 | 0.653 | 0.04 | 0.045 |
| thick neck | x1 | 0.035 | 0.590 | 0.07 | 0.045 |
| medium | x1.5 | 0.119 | 1.378 | 0.08 | 0.112 |
| large | x2 | 0.283 | 2.453 | 0.11 | 0.241 |
| **Mushroom** |  |  |  |  |  |
| small | x0.67 | 0.080 | 1.140 | 0.07 | 0.081 |
| medium | x1 | 0.271 | 2.567 | 0.10 | 0.232 |
| thin neck | x1 | 0.270 | 2.689 | 0.08 | 0.232 |
| thick neck | x1 | 0.272 | 2.507 | 0.13 | 0.232 |
| large | x1.33 | 0.643 | 4.568 | 0.13 | 0.526 |
| **filopodia-shaped** |  |  |  |  |  |
| small | x0.5 | 0.017 | 0.717 | 0.05 | 0.031 |
| medium | x0.75 | 0.058 | 1.609 | 0.08 | 0.064 |
| large | x1 | 0.138 | 2.860 | 0.10 | 0.127 |

**Table S2:** A list of spine apparatus variations.

| **Geometry** | **SA size** | **SA Volume ($\mu\text{m}^3$)** | **Cytoplasm Volume ($\mu\text{m}^3$)** |
| --- | --- | --- | --- |
| Thin | Small | 0.00211 | 0.033 |
|  | Medium | 0.00465 | 0.030 |
|  | Large | 0.00867 | 0.026 |
| Mushroom | Small | 0.0160 | 0.255 |
|  | Medium | 0.0358 | 0.235 |
|  | Large | 0.0676 | 0.203 |

**Table S3:** Table of values for realistic geometries.

| Spine Number | Shape | Volume ($\mu\text{m}^3$) | Surface Area ($\mu\text{m}^2$) | PSD Area ($\mu\text{m}^2$) |
| --- | --- | --- | --- | --- |
| 13 | Mushroom | 0.157 | 2.457 | 0.26 |
| 17 | Filopodia | 0.091 | 1.916 | 0.06 |
| 18 | Mushroom | 0.243 | 3.383 | 0.14 |
| 37 | Filopodia | 0.075 | 1.756 | 0.03 |
| 39 | Thin | 0.045 | 1.078 | 0.04 |
| 41 | Thin | 0.091 | 1.710 | 0.07 |

To graphically solidify the relationship between the calcium and the various synaptic weight update terms, we use artificial calcium transients as input into the synaptic weight model (Figure S2a). We consider three different cases, 1. a high calcium level (1000 ions) for 5 ms, 2. a lower calcium level (300 ions) for 5 ms, and 3. a high calcium level (1000 ions) for 10 ms. We can clearly see that for the prescribed calcium inputs, the *τ_w_* learning rate filters for rate of change to synaptic weight during the calcium pulse, while Ω*_w_* determines if the synaptic weight increases or decreases (Figure S2b-c). We see that as the calcium transient moves within the prescribed thresholds of LTP and LTD, the learning rate increases and decreases, while the Ω*_w_* term flips between negative, positive, and no change. For the selected LTP and LTD thresholds, both high calcium level cases lead to synaptic strengthening while the lower calcium level case leads to synaptic weakening (Figure S2d).

#### S3.2 Simulation results versus other geometric parameters show various trends

We plot max Ca^2+^ peak, decay time constant, and synaptic weight against volume for all size variations of filopodia-shaped spines, thin spines, mushroom spines, and mushroom spines with spine apparatus Figure S3. We see similar trends across volume as we observe across volume-to-surface area ratio. To understand the effect of volume on these calcium readouts, we plot several calcium transients for different geometries in terms of concentration, by dividing by the respective geometries’ volumes, Figure S4.

#### S3.3 Spine neck size shows differences in the large mushroom spines but not the smaller thin spines

The spine neck has long been discussed as a key parameter governing calcium signaling within dendritic spines (*38*). We also explored the effects of varying spine length and radius, while preserving spine volume. We first varied the spine neck on thin spines of the control volume, Figure S5a. We saw that while the calcium transients have considerable overlap, the thin-necked spine shows significant variance at later time points compared to the other spines, Figure S5b-c. We see no statistically significant differences between peak calcium values and only decay differences between the thinnest and thickest necks, Figure S5d-e. Synaptic weight changes for the thin spines with different neck geometries showed no significant differences but were trended towards negative weight changes for thicker necks, Figure S5f. We next explored mushroom spines with thinner or thicker neck geometries but with the same volume as the mushroom control spine, Figure S6a. While the mean of the calcium transients appeared quite close, there was significant difference in variance for the mushroom spine with the thick neck, Figure S6b-c. We saw differences in peak calcium only between the thinnest and thickest of the mushroom neck cases, and no significant difference in decay time constant, Figure S6d-e. Synaptic weight calculations show that presence of the thinnest versus thickest neck on a mushroom spine does lead to statistically significant differences in synaptic weight updates, Fig. S6f. This indicates that spine neck morphology might have more implications for these larger mushroom spines, compared to the smaller thin spines.

#### S3.4 The presence of spine apparatus in thin spines cause no clear trend in synaptic weight update

We vary the size of spine apparatus in thin control spines with the spine apparatus acting as a calcium sink with SERCA pumps, Figure S7a. We see that the presence of spine apparatus makes the calcium transient response more complex with a double peak visible in the variance for thin spines, Figure S7b-c. While we can fit the peak calcium values and decay time constant trends against both volume (Figure S7d,e) and volume-to-surface area ratio (Figure S7g,h), spine apparatus presence shows no clear trend in synaptic weight change for thin spines and the differences were not statistically significant, Figure S7f.

#### S3.5 Comparison to previous experimental and computational results

We compare our calcium transients to previously reported experimental and computational studies, Figure S8. To more directly compare, we normalize the various readouts to 1 and time shift the traces to all begin around the same time point. We consider the mean transient of a small idealized thin spine (0.035 µm^3^) and large idealized mushroom spine (0.643 µm^3^) from our study. We see that the small thin spine decays quite quickly while the larger mushroom spine has decay dynamics more similar to previously published findings.

#### S3.6 Our previous deterministic results match the qualitative trends seen in these results

We previously published a deterministic reaction diffusion model of calcium dynamics in dendritic spines of different morphologies (*35*). We found trends in the peak calcium concentration over spine volumes in that work and wanted to directly compare those results to our findings in this work. Using the results from (*35*), we integrate calcium concentration over the spine volume at each time point and find the peak calcium in ions and fit the decay dynamics of the calcium transient with an exponential decay function, *c* · exp(−*kt*). We compare the peaks and decay time constants over both volume and volume-to-surface area ratio, and find the same qualitative trends as our findings in this current work, Figure S9. We convert our current findings into concentration by dividing the total calcium ion transients by the geometry volumes, and consider five example transients for the three control idealized geometries and three realistic geometries, Figure S4. We find that the thin spines have the highest concentrations, followed by filopodia, and then mushroom spines.

#### S3.7 Synaptic weight changes depends on calculations with ions versus concentration

Synaptic weight update equations are typically phenomenological relationships based on Ca^2+^. Historically, many mathematical models considering synaptic weight changes have considered synaptic weight changes in terms of concentration (*10, 12, 18*). In this model, we consider Ca^2+^ in terms of Ca^2+^ ions. We want to consider if the use of ions versus concentration influences the synaptic weight update results. We converted the synaptic weight equations by converting the parameters from units involving molecules to concentration by dividing by the average spine volume (~0.06 µm^3^) and converting to µM. We convert all the Ca^2+^ transients to µM by dividing by each respective spine geometry volume and modifying units. We plot the synaptic weight change at 35 ms for all simulations when considering ions versus concentration Figure S10. We see that synaptic weight change predictions do change when using ions versus concentration because the concentration also considers the volume of the spines. Using concentration leads to a decreasing trend in synaptic weight with increasing volume which is the opposite of the trend seen using ions. We do however still see protrusion-type specific trends within the overall dynamics. There are several considerations to make during this comparison. First, as mentioned, the synaptic weight equations used are phenomenological relationships between Ca^2+^ and the concept of synaptic weight which captures the idea of synaptic strengthening which would actually occur through the insertion of receptors, such as AMPAR, and potentially spine volume increase. It remains unclear if total ion count, which is a global consideration of the whole spine, or Ca^2+^ concentration, which considers the local environment, is the correct value to consider for synaptic weight calculations. Furthermore, we used average concentration in Figure S10c-d) but dendritic spines are known to have signaling nanodomains, so it could be possible that it would be more accurate to consider peak concentration instead of average concentration for this calculation. Additionally, it is possible that the thresholds for LTP versus LTD need to be modified when considering a global reading, such as total ions in the spine, versus a local measurement, such as local concentration. Should synaptic weight change depend on the total amount of Ca^2+^ influx or the local environment within the spine? This is an ongoing consideration that needs further analysis and discussion.

Another consideration with this analysis is that the synaptic weight update is calcium-dependent. We plot the dynamics of the filopodia-shaped spine for a more direct comparison between calcium transients in terms of calcium ion and concentration, Figure S11. We can see that even through there is more calcium ion influx as the spines get larger, that increase in influx is not proportional to the increase in spine volume. Said another way, calcium influx is sublinear with respect to volume increase so larger spines have lower calcium transients. This is a consequence of our assumption of constant receptor density. This idea has been explored quite elegantly in (*12*). Therefore calcium influx relate to spine volume should continue to be explored and the consequences of geometry-dependent calcium trends on phenomenological relationships for synaptic weight need to be investigated further.

#### S3.8 Two-tailed ***t***-test results for all stochastic simulations

We conduct two-tailed *t*-test calculations between all stochastic simulations for both idealized and real geometries for max Ca^2+^ peak, decay time constant, and synaptic weight change. We display both the h and p-value for each comparison, Figure S12. We use a p threshold of 0.05 to determine the binary h value. A p-value smaller than 0.05 indicates that the two results are statistically different and produce a h-value of 1. Reversely, a p-value larger than 0.05 indicates that the two results are not statistically different and produce a h-value of 0. p-values have been truncated at two decimal points.

#### S3.9 Synaptic weight predictions from a single calcium transient mimic those from a pulse train

For all simulations so far, synaptic weight prediction has used a single calcium transient as input. However, usually a dendritic spine receives a pulse train of activation, so we want to consider the consequences of a pulse train of calcium transients on synaptic weight updates. To do this, for each 50 calcium transients per geometry, we repeated the calcium temporal dynamics 10 times with 35 ms between the initiation of the calcium spikes. This induction protocol of ~ 29 Hz is within the regime expected to begin to produce LTP in dendritic spines (*97, 98*). We compare the synaptic weight update for a single pulse to the update for a pulse train, Figure S13a vs b. We see that synaptic weight change increases in magnitude for the pulse train condition but keeps the same general trends observed in the single pulse simulations, indicating the a single pulse does give a good approximation of pulse train dynamics at this frequency. However, it is clear that the conversion between single pulse and pulse train is not as clear for the realistic geometry cases. Additionally, the pulse train predictions showed a increase in standard deviation, with more cases showing the ability to increase or decrease their synaptic weight. Therefore, while the mean synaptic weight change for each condition held across a single pulse and multiple pulses, the pulse train widens the synaptic weight regime that a spine can enter.

**Figure S2:**
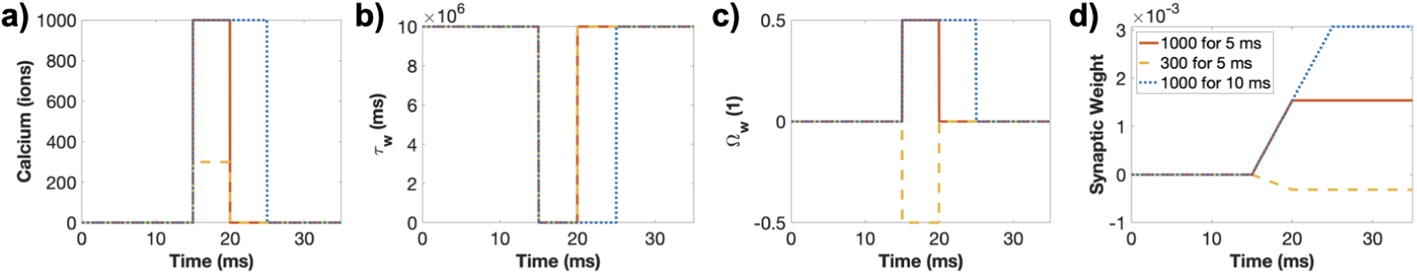
Artificial calcium transients demonstrate how learning rate, Ω*_w_*, and synaptic weight depend on calcium temporal dynamics. To illustrate the relationship between calcium (a), *τ_w_* (b), Ω*_w_* (c), and synaptic weight (d), we construct three artificial calcium profile with heaviside functions. The calcium profiles are 1. 1000 ions for 5 ms (red line), 2. 300 ions for 5 ms (yellow dashed line), and 3. 1000 ions for 10 ms (blue dotted line).

#### S3.10 Supplemental movies

##### S3.10.1 Supplemental Movie S1

**Sample movie of idealized filopodia simulation.** A single seed of an idealized filopodia simulation is shown for the whole time period from 0 to 35 ms. The plasma membrane mesh is shown in blue and the Ca^2+^ ions are red.

##### S3.10.2 Supplemental Movie S2

**Sample movie of idealized thin spine simulation.** A single seed of an idealized thin spine simulation is shown for the whole time period from 0 to 35 ms. The plasma membrane mesh is shown in blue and the Ca^2+^ ions are red.

##### S3.10.3 Supplemental Movie S3

**Sample movie of idealized mushroom spine simulation.** A single seed of an idealized mushroom spine simulation is shown for the whole time period from 0 to 35 ms. The plasma membrane mesh is shown in blue and the Ca^2+^ ions are red.

**Figure S3:**
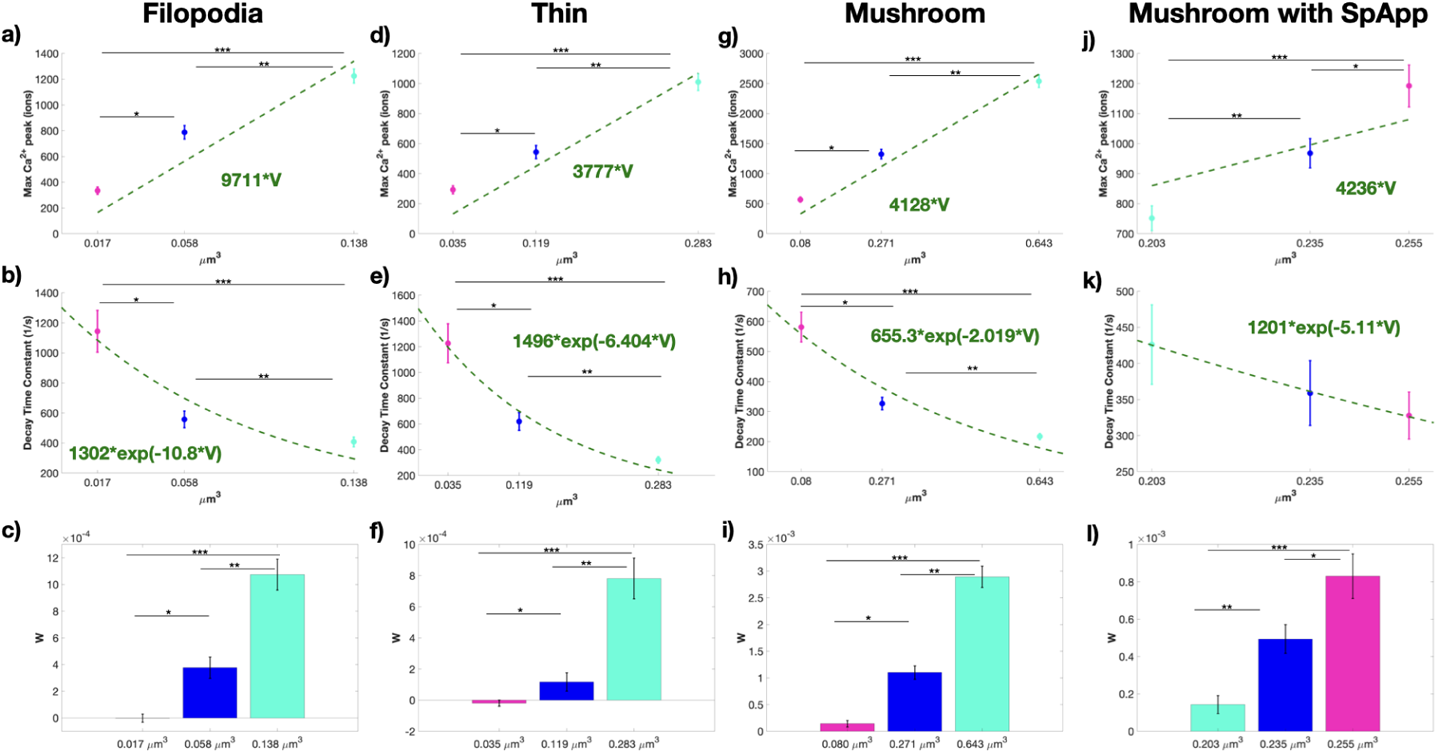
Trends across volume are similar to trends across volume-to-surface area ratio. Peak calcium levels, decay time constant, and synaptic weight updates for size variations given as volumes for filopodia-shaped spines (a-c), thin spines (d-f), mushroom spines (g-i), and mushroom spines with spine apparatus (j-l). Peak calcium is fit with a line with a fixed zero intercept.

**Figure S4:**
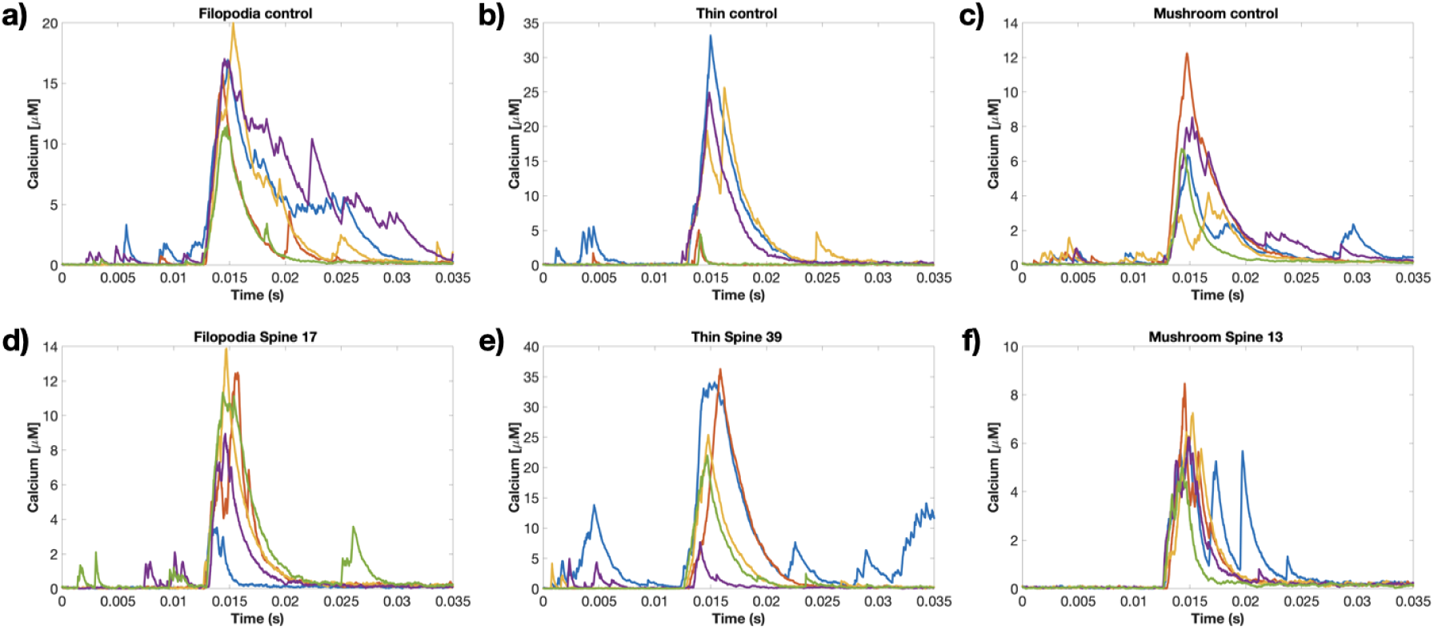
Examples of calcium transients in terms of concentration for various geometries. Five examples of calcium transients in terms of concentration for the filopodia control geometry (a), thin control geometry (b), mushroom control geometry (c), realistic filopodia spine 17 (d), realistic thin spine 39 (e), and realistic mushroom spine 13 (f).

**Figure S5:**
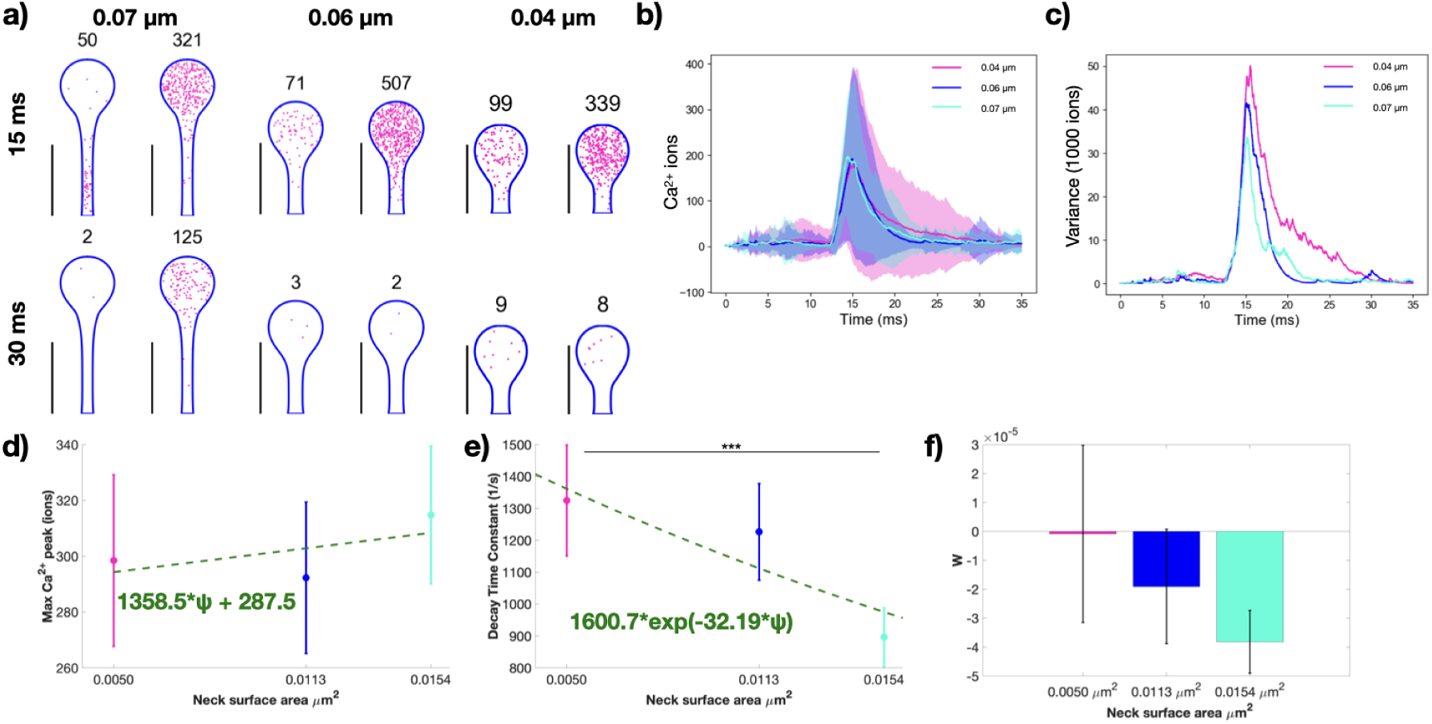
Effect of spine neck variation on synaptic plasticity in thin spines. a) Spatial plots at 15 and 30 ms for thin spines of the same volume with different neck geometries (neck radius of 0.04, 0.06, 0.07 *µm*). The number above each spine corresponds to the number of calcium ions present at that time point. Scale bar: 2 µm. Calcium ions over time (b) and variance, displayed as variance divided by 1000 ions (c), for all three thin spines with different neck cases. Shaded regions in (b) denote standard deviation. d) Peak calcium ion number for each thin spine with the mean and standard error (n=50) show no statistically significant differences using a two-tailed *t*-test. We fit the trend in peak calcium as a linear function of spine neck base surface area; *r*^2^ = 0.0009 for the linear fit. e) We fit the decay portion of each calcium transient with the exponential decay function *c* · exp(−*kt*). The decay time constant mean and standard error (n=50), *k*, only shows statistically significant differences between the thin and thick necks; p*** = 0.0322 from a two-tailed *t*-test. We fit the trend in decay time constants as a function of spine neck base surface area with an exponential *a* · exp(−*bψ*), where *ψ* is the spine neck base surface area; *r*^2^ = 0.0256 for the exponential fit. f) Calculated synaptic weight change at the last time point for all three thin spines shows no statistically significant difference due to neck size.

**Figure S6:**
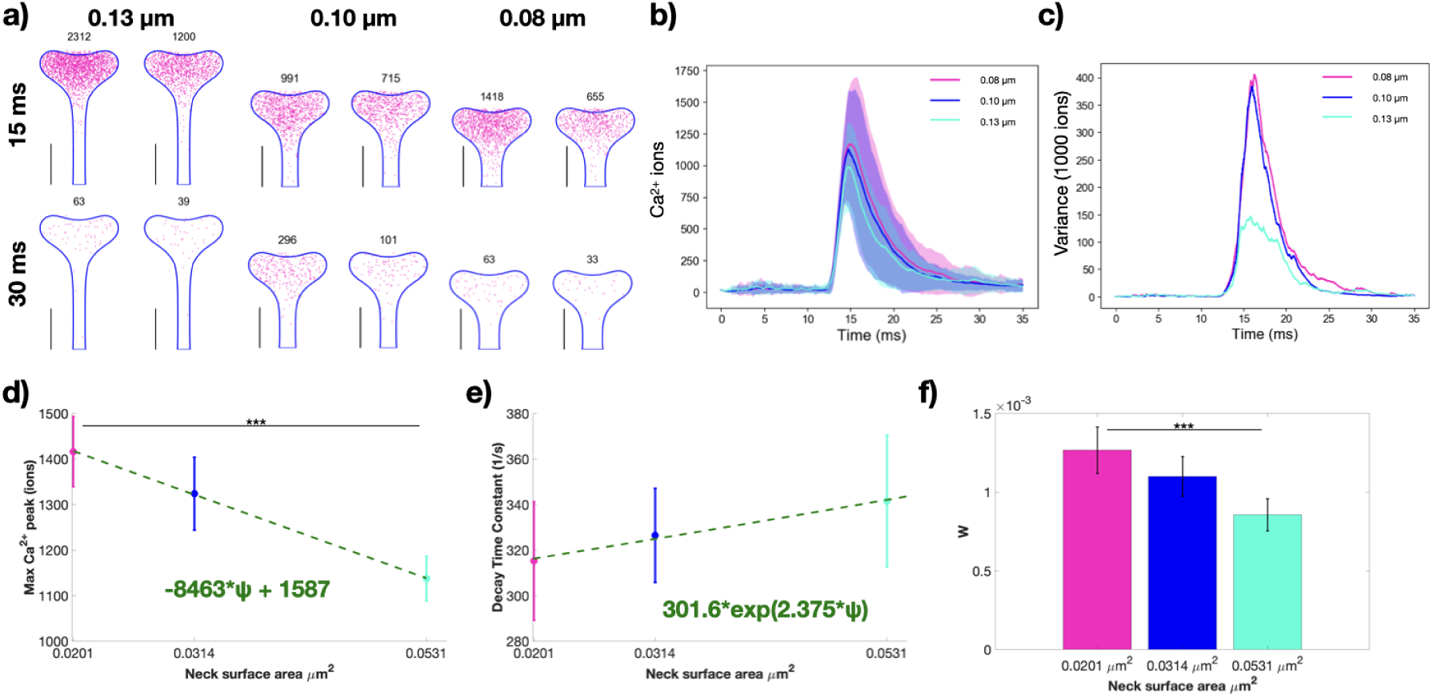
Effect of spine neck variation on synaptic plasticity in mushroom spines. a) Spatial plots at 15 and 30 ms for mushroom spines of the same volume with different neck geometries (neck radius of 0.08, 0.10, 0.13 *µm*). The number above each spine corresponds to the number of calcium ions present at that time point. Scale bar: 2 µm. Calcium ions over time (b) and variance, displayed as variance divided by 1000 ions (c), for all three mushroom spines with different neck cases. Shaded regions in (b) denote standard deviation. d) Peak calcium ion number for each mushroom spine with the mean and standard error (n=50) show statistically significant differences between the thin and thick spines; p*** = 0.0029 using a two-tailed *t*-test. We fit the trend in peak calcium as a linear function of spine neck base surface area; *r*^2^ = 0.0528 for the linear fit. e) We fit the decay portion of each calcium transient with the exponential decay function *c* exp(−*kt*). The decay time constant mean and standard error (n=50), *k*, shows no statistically significant differences from a two-tailed *t*-test. We fit the trend in decay time constants as a function of spine neck base surface area with an exponential *a* exp(−*bψ*), where *ψ* is the spine neck base surface area; *r*^2^ = 0.0036 for the exponential fit. f) Calculated synaptic weight change at the last time point for all three mushroom spines only shows a statistically significant difference between the thin and thick spines, p*** = 0.0244 from two-tailed *t*-test.

**Figure S7:**
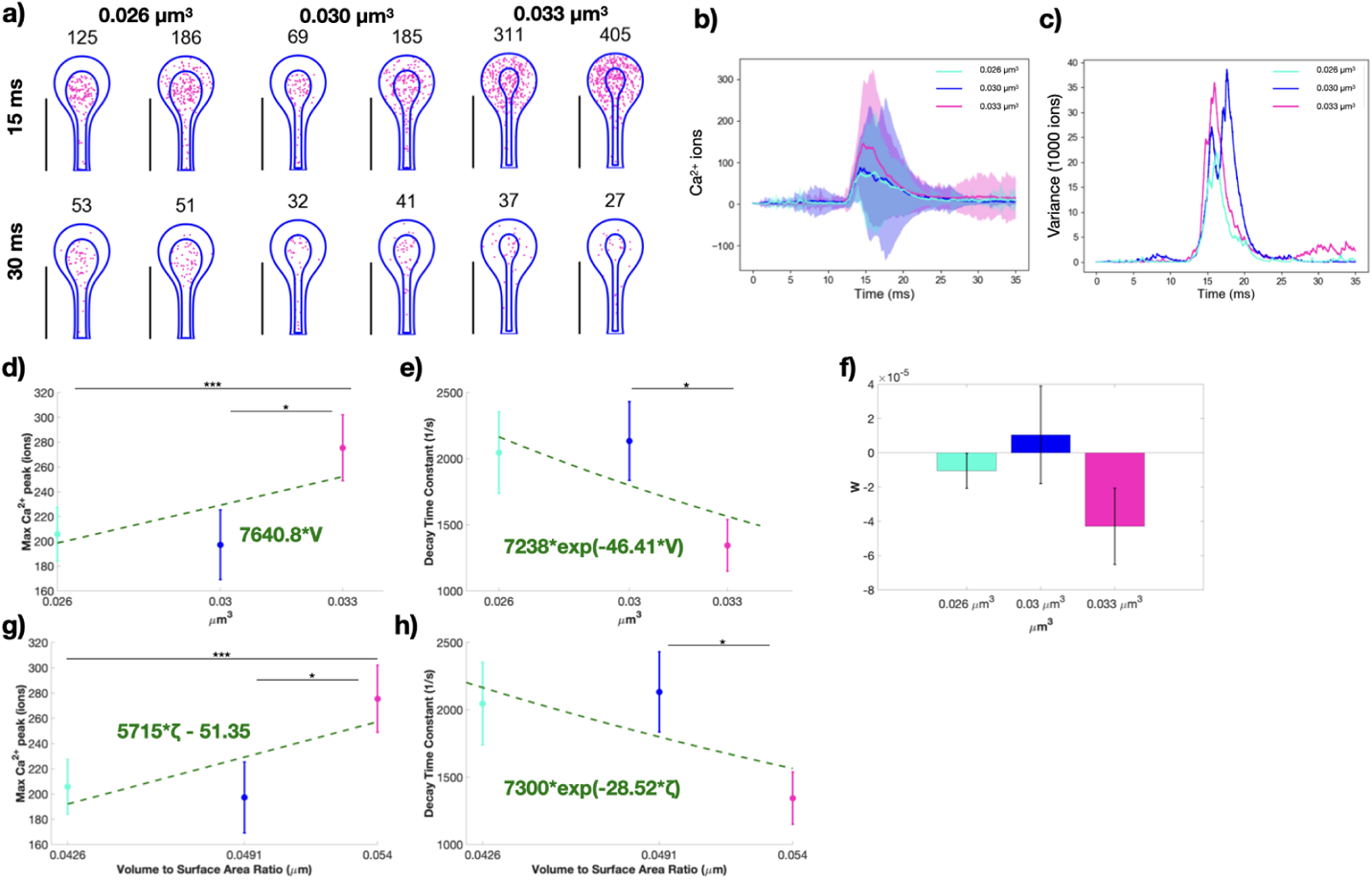
Spine apparatus size modulates synaptic weight change in thin spines. a) Spatial plots at 15 and 30 ms for thin spines with spine apparatus of different volumes (spine cytosolic volumes of 0.026, 0.030, 0.0.033 *µm*^3^). The numbers on top of the shape indicate the total number of calcium ions at that instant in both the spine apparatus and cytoplasm. Calcium ions over time as mean and standard deviation (b) and variance, displayed as variance divided by 1000 ions (c), for all three thin spines with different spine apparatus sizes. Shaded regions in (b) denote standard deviation. d) Peak calcium ion number for each thin spine with a spine apparatus, with the mean and standard error (n=50), show statistically significant differences between two of the three paired cases; p* = 0.0461; p*** = 0.0453 from two-tailed *t*-test. We fit the trend in peak values with a linear function against the cytoplasm volume; *r*^2^ = 0.0145 for the linear fit. e) We fit the decay dynamics of each calcium transient with *c* · exp(−*kt*) and report the decay time constant, k, as a mean and standard error (n = 50). We find only find statistically significant differences between the second and third spines; p* = 0.0289 from a two-tailed *t*-test. We fit the trend in decay time constants as a function of cytosolic volume with an exponential *a* · exp(−*bV*), where V is the cytosolic volume; *r*^2^ = 0.0177 for the fit. f) Calculated synaptic weight change at the last time point for all three thin spines shows no statistically significant difference due to spine apparatus size. We also plot peak calcium ion number and decay time constant against the cytosolic volume to surface area ratio, g and h, respectively. g) We fit the trend in peak values with a linear function against the volume-to-surface area ratio; *r*^2^ = 0.0214 for the linear fit. h)We fit the trend in decay time constants as a function of volume-to-surface area ratio with an exponential *a* · exp(−*−bζ*), where *ζ* is the volume-to-surface area ratio; *r*^2^ = 0.0178 for the fit.

**Figure S8:**
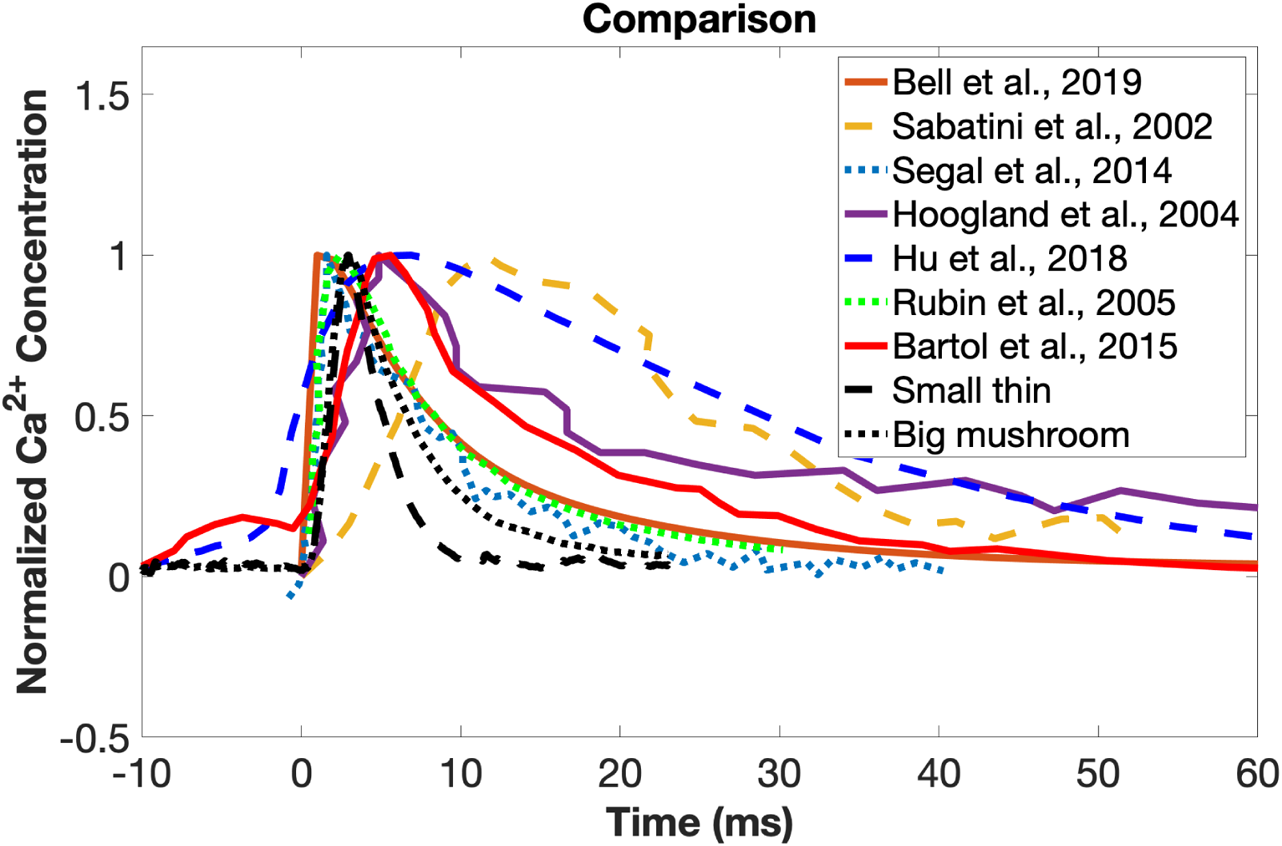
Normalized calcium transients from different experimental and computational studies. We plot the temporal dynamics of the small idealized thin spine (0.035 µm^3^) and large idealized mushroom spine (0.643 µm^3^) versus reported experimental calcium transients from previous studies (*64–66*) and computational model results from previous studies (*35, 37, 62, 63*). The various experimental transients are reported in terms of fluorescence, and we assume the transients are linearly proportional to concentration (*110*). We normalize the various transients and time shift them for a more direct comparison. We plot Fig. 1 F from Sabatini et al. (2002) (*64*), Fig. 1 from Segal and Korkotian (2014) (*66*), and Fig. 2 D from Hoogland and Saggau (2004) (*65*). We are also comparing Fig 3 of Bell et al (2019) (*35*), Fig. 5 of Hu et al. (2018) (*63*), Fig. 1 A of Rubin et al. (2005) (*62*), and Fig. 7 I of Bartol et al. (2015) (*37*).

**Figure S9:**
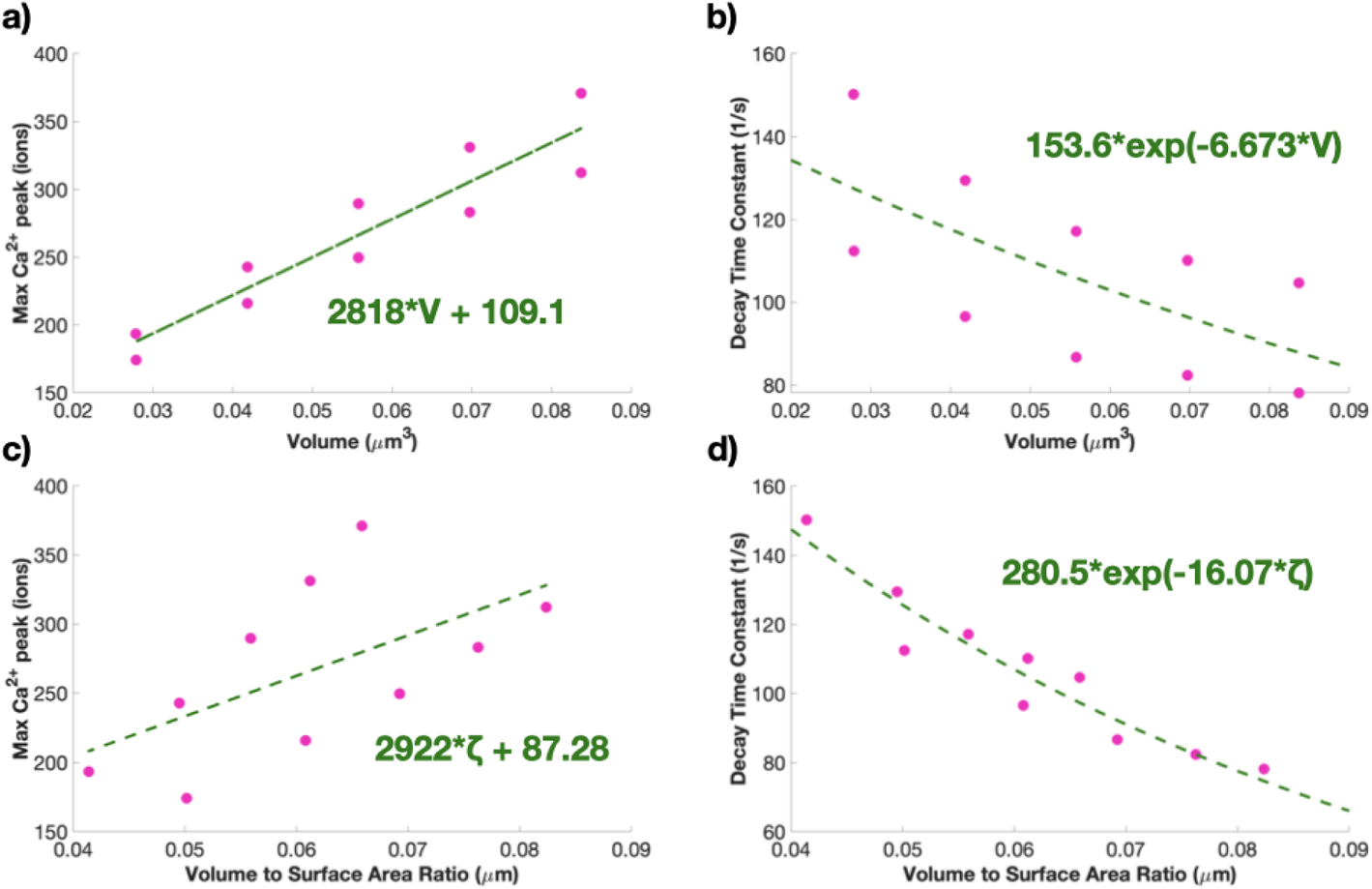
Previous calcium simulation results match the qualitative trends in these results. a) We fit the trend in peak values with a linear function against the cytoplasm volume; *r*^2^ = 0.8776 for the linear fit. We fix the y intercept at zero. b) We fit the decay dynamics of each calcium transient with *c* · exp(−*kt*) and report the decay time constant, k. We fit the trend in decay time constants as a function of cytosolic volume with an exponential *a* · exp(−*bV*), where V is the cytosolic volume; *r*^2^ = 0.4283 for the fit. c) We fit the trend in peak values with a linear function against the volume-to-surface area ratio; *r*^2^ = 0.3492 for the linear fit. h) We fit the trend in decay time constants as a function of volume-to-surface area ratio with an exponential *a* · exp(−*bζ*), where *ζ* is the volume-to-surface area ratio; *r*^2^ = 0.9054 for the fit.

**Figure S10:**
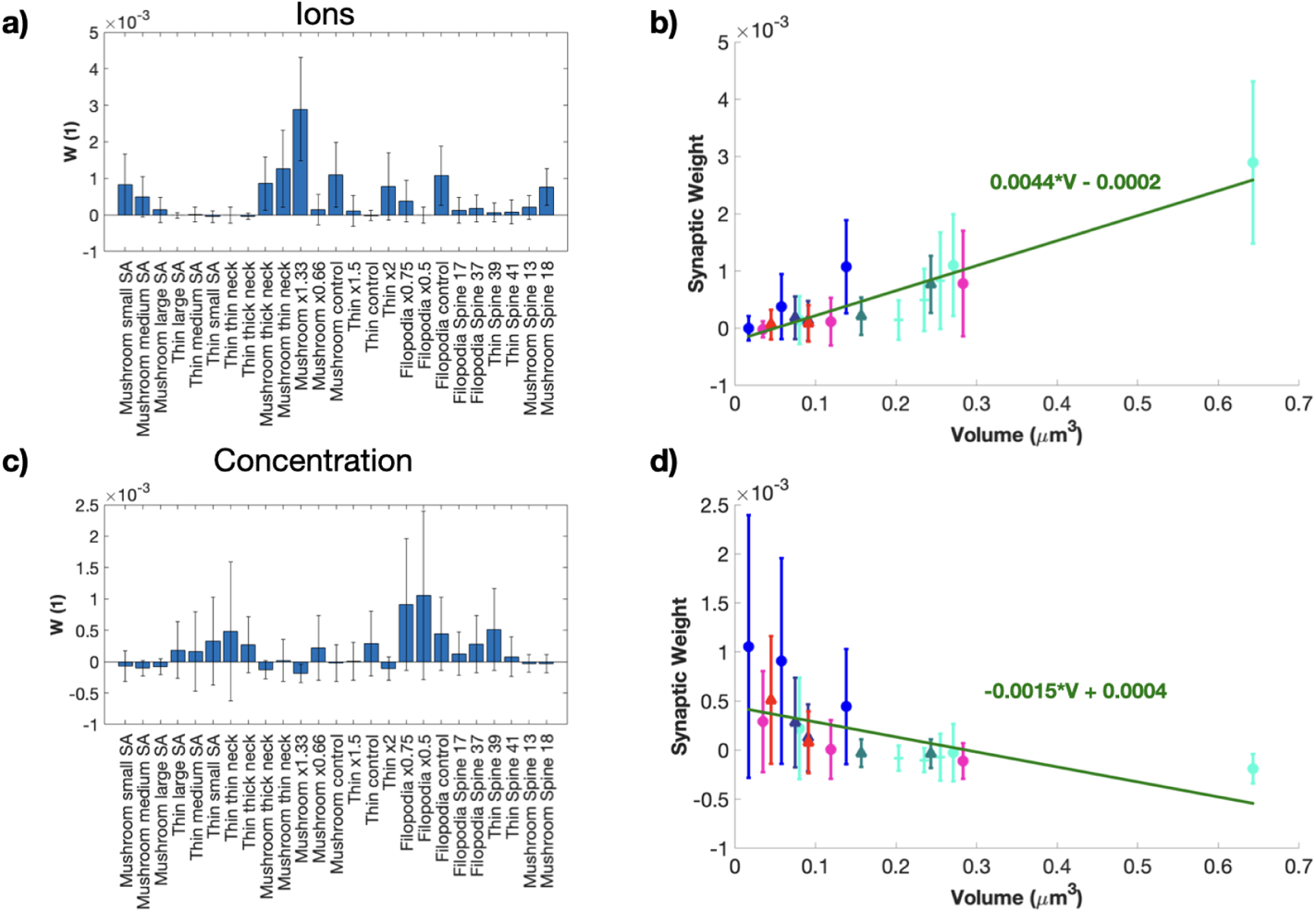
Synaptic weight updates when considering Ca^2+^ in terms of ions or concentration. Synaptic weight updates for each stochastic idealized and real geometry simulation when synaptic weight calculations are in terms of ions (a-b) and concentration (c-d). We plot the synaptic weight changes against the spine volume for calculations using ions (b) and concentration (d). We fit the trends using a linear function of volume. We get *r*^2^ = 0.4635 for the ion fit and *r*^2^ = 0.1229 for the concentration fit.

**Figure S11:**
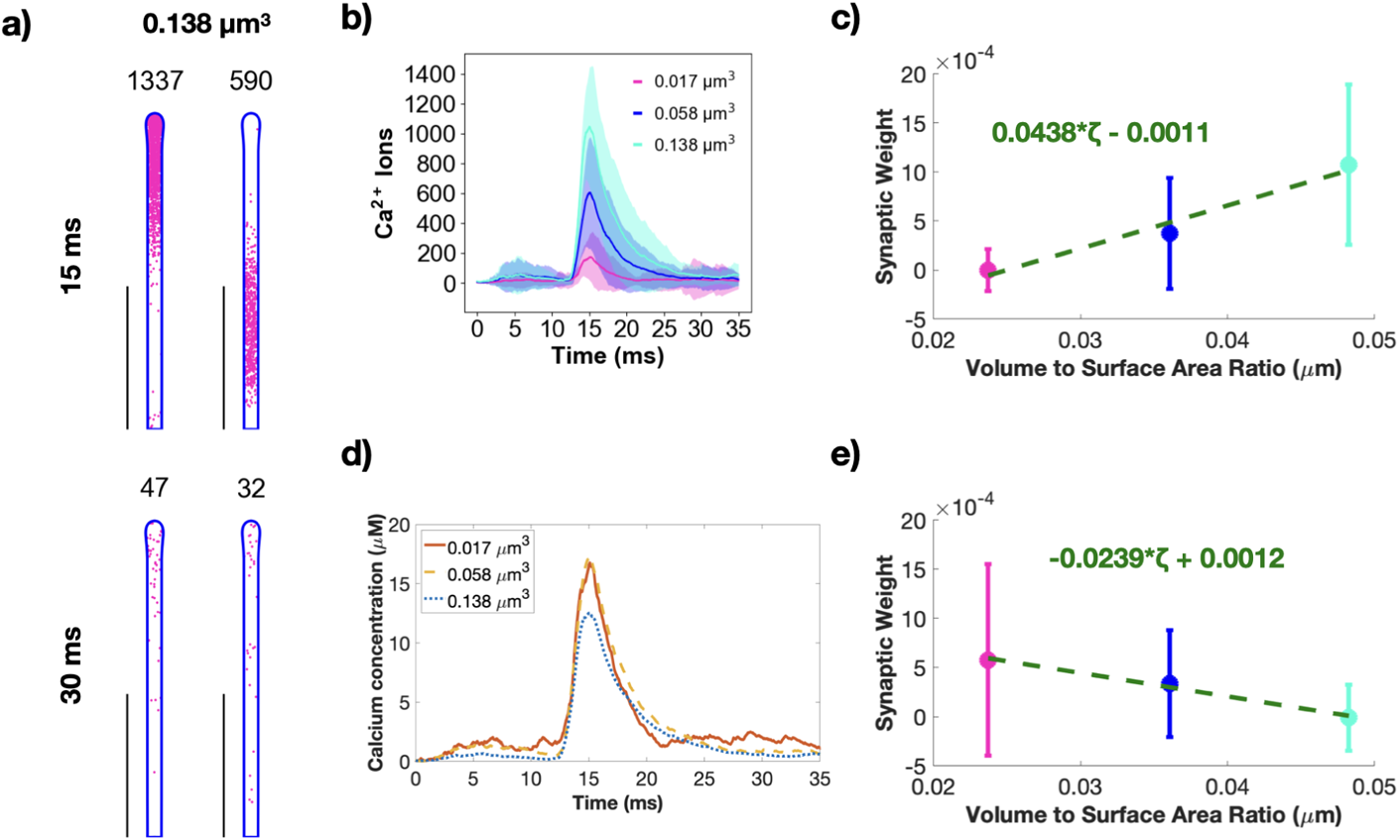
Synaptic weight updates when considering Ca^2+^ in terms of ions or concentration for the filopodia-shaped spines. a) Spatial plots illustrating Ca^2+^ localization at 15 and 30 ms for a filopodia-shaped spine with volume 0.138 µm^3^. The number above each geometry corresponds to the number of Ca^2+^ in that frame. Two random seeds are shown. Scale bars: 2 µm. b) Mean (solid) and standard deviation (shaded area) of Ca^2+^ transients across 50 simulations for each of the three filopodia-shaped spine sizes (0.017, 0.058 and 0.138 µm^3^). c) Synaptic weight prediction for each of the filopodia geometries calculated as a function of total calcium ions. We fit the trends using a linear function of volume to surface area ratio. We get *r*^2^ = 0.3594 for the ion fit. d) Mean of the calcium transients for each filopodia-shaped spine sizes converted to concentration. e) Synaptic weight prediction for each of the filopodia geometries calculated as a function of calcium concentration. We fit the trends using a linear function of volume to surface area ratio. We get *r*^2^ = 0.1143 for the concentration fit. We find statistically significant differences between the first and third spines, and between the second and third spines; p_12_ = 0.1301; p_23_ = 2.2567 × 10^−4^; p_13_ = 1.1347 × 10^−4^ from two-tailed *t*-test, where 1, 2, and 3 correspond to the spines in increasing volume.

**Figure S12:**
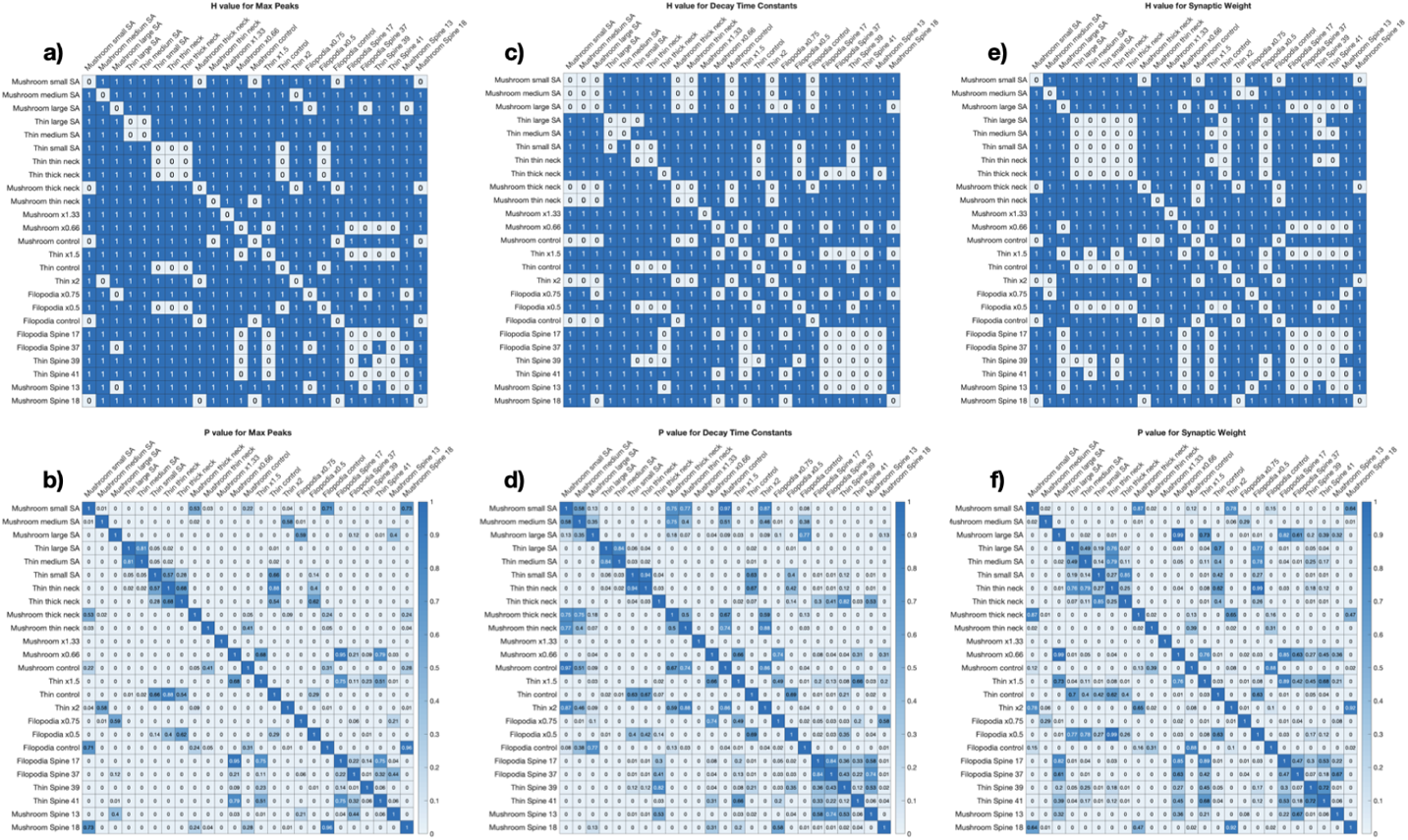
Two-tailed *t*-test comparison between all simulations. We conduct two-tailed *t*-test between all simulations and display the h value and p-value for max Ca^2+^ peaks (a-b), decay rate constant (c-d), and synaptic weight change (e-f). Displayed p-values are truncated at two decimal points.

**Figure S13:**
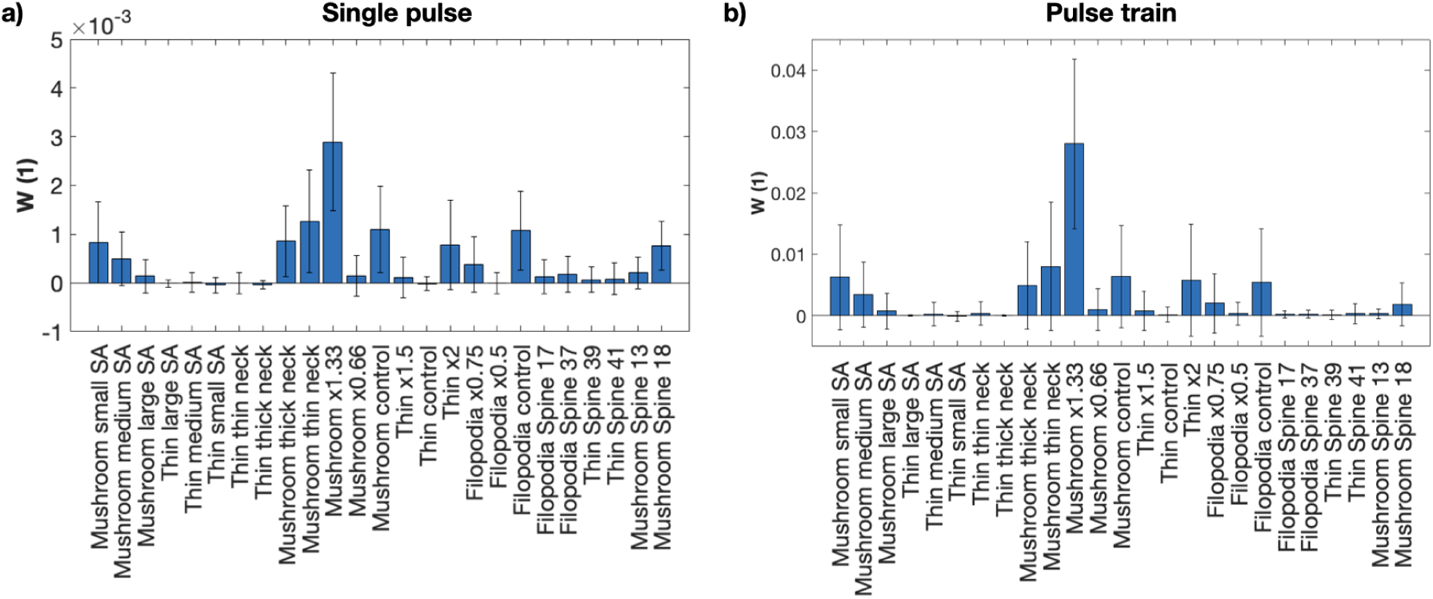
Single pulses and multiple pulses show similar trends in synaptic weight updates across different geometries. a) Synaptic weight change due to a single pulse of calcium for the different spine geometries. b) Synaptic weight change due to multiple pulses of calcium for the different spine geometries.

##### S3.10.4 Supplemental Movie S4

**Sample movie of realistic mushroom spine 13 simulation.** A single seed of a realistic mushroom spine 13 simulation is shown for the whole time period from 0 to 35 ms. The plasma membrane mesh is shown in blue and the Ca^2+^ ions are red.

##### S3.10.5 Supplemental Movie S5

**Sample movie of realistic filopodia 17 simulation.** A single seed of a realistic filopodia 17 simulation is shown for the whole time period from 0 to 35 ms. The plasma membrane mesh is shown in blue and the Ca^2+^ ions are red.

##### S3.10.6 Supplemental Movie S6

**Sample movie of realistic mushroom spine 18 simulation.** A single seed of a realistic mushroom spine 18 simulation is shown for the whole time period from 0 to 35 ms. The plasma membrane mesh is shown in blue and the Ca^2+^ ions are red.

##### S3.10.7 Supplemental Movie S7

**Sample movie of realistic filopodia 37 simulation.** A single seed of a realistic filopodia 37 simulation is shown for the whole time period from 0 to 35 ms. The plasma membrane mesh is shown in blue and the Ca^2+^ ions are red.

##### S3.10.8 Supplemental Movie S8

**Sample movie of realistic thin spine 39 simulation.** A single seed of a realistic thin spine 39 simulation is shown for the whole time period from 0 to 35 ms. The plasma membrane mesh is shown in blue and the Ca^2+^ ions are red.

##### S3.10.9 Supplemental Movie S9

**Sample movie of realistic thin spine 41 simulation.** A single seed of a realistic thin spine 41 simulation is shown for the whole time period from 0 to 35 ms. The plasma membrane mesh is shown in blue and the Ca^2+^ ions are red.

##### S3.10.10 Supplemental Movie S10

**Sample movie of idealized mushroom spine with a spine apparatus simulation.** A single seed of an idealized mushroom spine with a spine apparatus simulation is shown for the whole time period from 0 to 35 ms. The plasma membrane and spine apparatus membrane mesh are shown in blue and the Ca^2+^ ions are red.

